# The antiviral Interferon pathway drives astrocyte aging and motor decline

**DOI:** 10.1101/2025.11.12.688147

**Authors:** Lara Labarta-Bajo, Ayesha R Thanawalla, Irene L Gutierrez, Su May Lei Soe, Brent Y Chick, Ioanna Andritsogianni, Setareh Metanat, Diana C Hargreaves, Susan M Kaech, Eiman Azim, Nicola J Allen

## Abstract

Aging encompasses low-level inflammation and motor decline. Astrocytes are neuroregulatory glial cells that change in aging, particularly in the cerebellum, which is essential for movement coordination. Regulation and functionality of cerebellar astrocytes in aging is unknown. We show that antiviral type I Interferons (IFN-I) drive motor deficits and regional astrocyte aging. Transcriptomics reveal that cerebellar astrocytes, but not cortical, exhibit an antiviral state that intensifies with age, with increased expression of Stat1. Aged mice display motor deficits similar to humans that improve after peripheral IFN-I receptor neutralization, whereas astrocyte Stat1 induces motor deficits during chronic inflammation in adults. While strong systemic inflammation induces astrocyte antiviral state, in aging, chromatin de-repression of Stat1 and nucleotide sensors in cerebellar astrocytes amplifies local IFN-I signaling. We identify functional interaction between a classical immune pathway and astrocytes, representing an actionable strategy to preserve motor function in aging.

## Introduction

Human aging is characterized by a rise in inflammatory pathways in peripheral organs and within the central nervous system (CNS)^1–4^. Progressive decrease in motor function is also a characteristic of healthy aged human individuals, a trait that precedes and predicts cognitive decline^5–7^. In fact, the cerebellum, a brain region that regulates motor coordination, posture and balance^8,9^, accumulates age-related molecular and functional changes earlier than most other brain regions in mice and humans^3,10,11^. While inflammatory pathways associate with advanced age and functional decline, little is known about the origins and causative role of inflammation in age-related molecular changes leading to motor decline.

Astrocytes are glial cells that play key neuroregulatory roles during development, and in adulthood^12^. Similar to neurons, astrocytes in the developed CNS are regionally heterogeneous, suggestive of specialization^3^. Within the CNS, astrocytes are among the cells that accumulate the largest number of transcriptional changes in advanced age^3^, however their regulation and functionality in aging is not well understood. Astrocytes maintain regional heterogeneity in aging, and accumulate changes in a region-specific manner, with a larger magnitude of changes in the cerebellum, compared to hypothalamus, cortex, striatum or hippocampus^13,14^. Given the role of the cerebellum in motor coordination, this suggests that regional properties of astrocytes are relevant in age-related motor decline.

Aging constitutes a state of chronic, low-level inflammation involving the activation of innate immune pathways involved in anti-viral defense, including those induced by Interferons, TNFα, IL-1, and IL-6^1,3,4,15–17^. High-level inflammation induces sickness and influences motor skill^18–20^, and exercise promotes immune changes, including type I Interferon (IFN) signaling^21,22^, which promotes anti-viral responses after infection^23^. The receptor for type I IFN cytokines is broadly expressed across the body, and in most CNS cell types including in astrocytes^3,24^. Thus, antiviral gene up-regulation in aging implicates the type I IFN pathway, which promotes cognitive deficits^25,26^, but whose role in astrocytes and in motor decline is far less understood.

In this study we find that the type I IFN antiviral pathway promotes motor decline and astrocyte aging in the cerebellum. Transcriptional analysis establishes an antiviral state of cerebellar astrocytes, but not cortical, that is present in adulthood and intensifies with age, with expression of the transcription factor Stat1. Aged mice have motor deficits similar to humans, that are alleviated after systemic type I IFN signaling is neutralized, and in adult mice, astrocyte Stat1 constitutes a vulnerability, leading to motor deficits during chronic inflammation. Low-level peripheral inflammation does not remarkably impact astrocyte state, while high-level systemic inflammation induces astrocyte antiviral state in both the cortex and cerebellum, and does not explain regionality. Instead, local IFN-I signaling promotes astrocyte antiviral signatures, and in aging, such antiviral state is amplified by increased chromatin accessibility leading to increased expression of antiviral gene Stat1, and double stranded RNA sensors, which can perpetrate antiviral responses. Overall, we uncover functional interaction between the type I IFN antiviral pathway and astrocytes, that can be targeted to alleviate astrocyte aging and motor deficits in the elderly.

## Results

### Astrocytes in the cerebellum exhibit an antiviral state that is exacerbated in aging

In aging, astrocytes in the cerebellum undergo more transcriptional changes than those in other brain regions^13,14^. To elucidate astrocyte regulation in the aged cerebellum we analyzed previous datasets that profiled actively translated mRNAs in astrocytes in the cortex, striatum, hippocampus, hypothalamus, and cerebellum of WT adult (2.5-4mo) and aged (20-24mo) mice^13,14^. Pathway analysis using differentially up-regulated genes in aged compared to adult astrocytes finds enrichment of numerous terms related to the antiviral response in the aged cerebellum, such as ‘response to virus’ or ‘regulation of the innate immune response’ (Fig. 1A, Table S1). Pathway-level enrichment of immunity-related genes is observed in astrocytes in the aged hippocampus and hypothalamus, albeit to a lesser extent, and is not detected in cortex or striatum (Fig. S1A-C & Table S1). We created a unified antiviral signature with genes involved in the response to neuroinflammation and with functional roles during infections, including in antigen presentation (e.g. *B2m*, *H2-d1*)^27^, RNA sensing (e.g. *Oas1a*, *Ifih1*)^28^, type I Interferon (IFN) production and antiviral gene expression (e.g. *Irf7*, *Stat1*)^29^ as well as negative regulators of the immune response (e.g. *Usp18*, *Trim30a*)^29,30^. All 58 genes are up-regulated in cerebellar astrocytes, and only a small number are up-regulated in other regions, thus showing regional enrichment in the cerebellum (Fig. 1B & S1D,E). Astrocytes up-regulate reactivity-associated genes during neuroinflammation^31–33^, and in aging across brain regions^13,14^. We find that a larger number of reactive genes are up-regulated in astrocytes in the cerebellum compared to other brain regions (Fig. 1B, S1F & Table S1)^13^. Antiviral genes are rapidly transcribed and translated in response to inflammation^29^. To elucidate if antiviral genes are actively induced, we analyzed bulk RNA-seq of astrocyte nuclei in the adult and aged cerebellum that was generated using Glyoxal Fixed Astrocyte Transcriptomics (GFAT)^34^ (Fig. S2A-D). This analysis shows up-regulation of antiviral and reactivity genes in aged (2yo) compared to adults (4mo), similar to changes observed using translatome analysis (Fig. 1C-D & S2H-I & Table S1). Thus, our analyses show persistent induction and translation of antiviral genes in astrocytes in the aged cerebellum.

**Figure 1.**
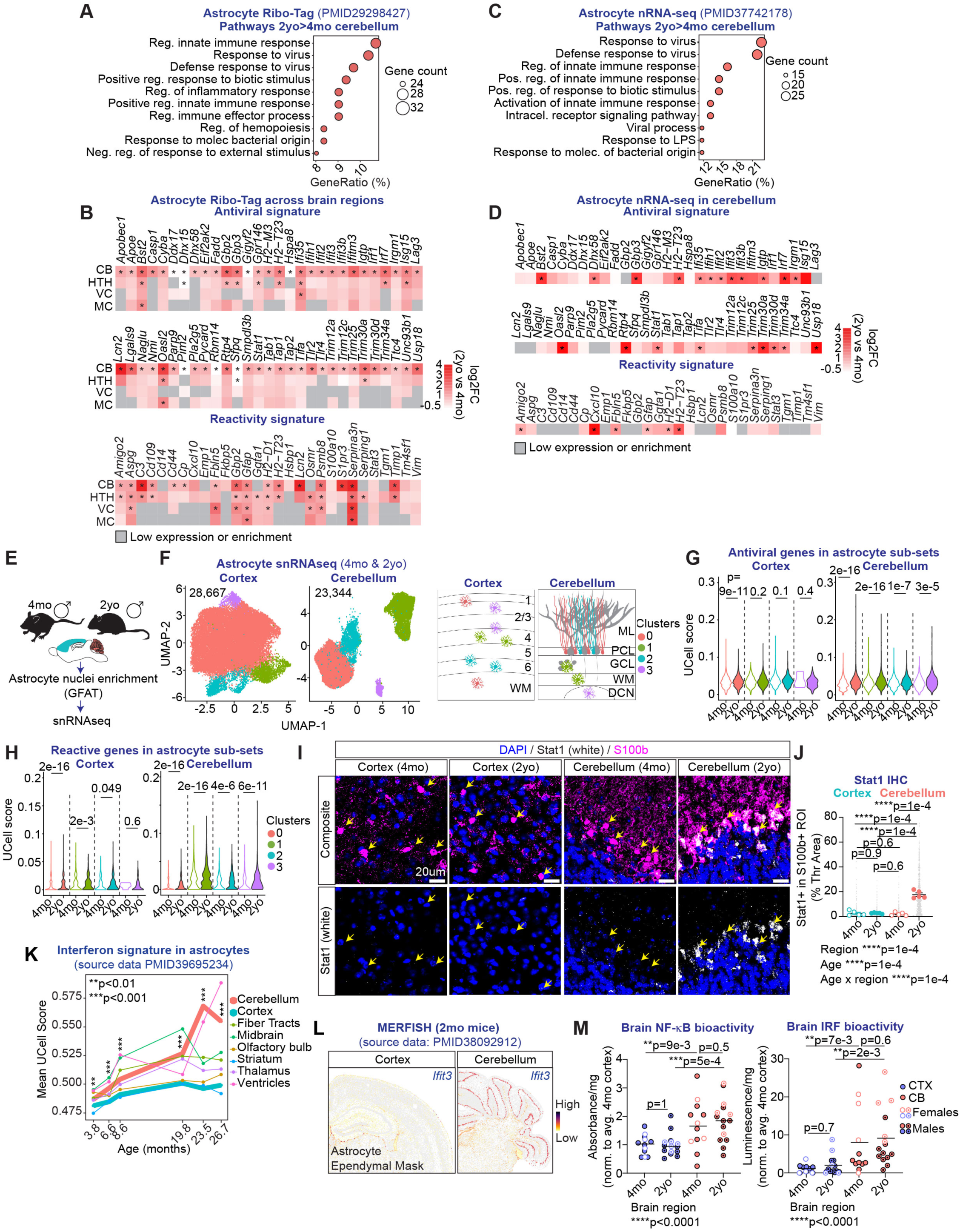
Astrocytes in the cerebellum exhibit an antiviral state that is exacerbated in aging. (A,C) Gene ontology (GO) pathway analysis was done using published astrocyte RNA-seq datasets that profiled astrocytes in the adult (4mo) and aged (2yo) cerebellum of WT male mice using Ribo-Tag (A) or nucleus (n)RNA-seq (C). Differentially-expressed genes (DEGs) in cerebellar astrocytes in 4mo vs 2yo mice are used with cut-offs: p.adj<0.05, absolute fold-change (FC)>1.5, enrichment in astrocytes over input>0.75, FPKM>1 (Ribo-Tag), and p.adj<0.05, FC>1.5, TPM>5 (nRNA-seq). Pathways with 10 or more DEGs overlapping with GO pathways are shown. GeneRatio refers to the percentage of DEGs, among all DEGs, that overlap with a GO pathway. (B,D) Heatmaps showing log2 fold change in gene expression of antiviral and reactivity-associated genes in astrocytes in the cerebellum (CB), hypothalamus (HTH), visual cortex (VC), and motor cortex (MC) profiled using Ribo-Tag (B), and in the cerebellum using nRNA-seq (D). Adjusted p-value<0.05*. (E-H) Single-nucleus (sn)RNA-seq of astrocyte nuclei in the cortex and cerebellum of 4mo and 2yo male WT mice. (F) Astrocyte sub-sets in the adult and aged cortex and cerebellum. Cortical sub-sets are located across cortical layers and in the white matter (WM), and cerebellar astrocyte sub-sets include Bergmann glia in the molecular layer and Purkinje Cell layers (ML, PCL), velate astrocytes in the granule cell layer (GCL), fibrous in the white matter (WM), and in the deep cerebellar nuclei (DCN). Numbers in the upper left corner indicate the number of astrocyte nuclei analyzed. (G,H) Antiviral and reactive signatures are scored in astrocyte sub-sets in the cortex and cerebellum using UCell. (I) Representative images show Stat1 protein in S100b-positive astrocytes in the aged (2yo) cerebellum, but not in cortex or adult (4mo) cerebellum. (J) Quantification of Stat1 signal overlapping with S100b+ cell soma in (I). Small grey dots are individual S100b+ soma, and circles are mouse averages. (K) Gene signature UCell scores for Interferon-related genes using a published dataset that profiles astrocytes across brain regions in adult and aged WT male mice. (L) Representative images show *Ifit3* mRNA expression in astrocytes and ependymal cells in the cerebellum, but not in cortex, of an adult (2mo) mouse. (M) Nuclear factor kappa-light-chain-enhancer of activated B cells (NF-kB) and Interferon regulatory factor (IRF) bioactivity in cortical and cerebellar brain lysates from 4mo and 2yo mice. Data are normalized to the average of the 4mo cortex. (B,D) Significance calculated using DESeq2, (G,H,K) Wilcox test, (J) 2-way ANOVA with Sidak’s multiple comparisons test, (M) Kruskal-Wallis with Dunn’s multiple comparison. (I,J) 5 mice/group, (M) 12-14 mice/group. (J,M) Horizontal bars indicate the mean.

Astrocytes are heterogeneous across and within brain regions^3,35^. To elucidate if antiviral programs are enriched in particular sub-sets, or rather, at the compartment level, we performed single-nucleus (sn) RNA-seq analysis of astrocytes in the cortex and cerebellum of adult (4mo) and aged (2yo) mice (Fig. 1E & S2E-G & S3). Cluster analysis of 28,677 cortical astrocytes shows up-regulation of reactivity-associated genes in most astrocytes in aging, but no increase in the expression of antiviral genes in any sub-set (Fig. 1F-H & S3A-D & Table S2). Thus, while astrocytes in the cortex change with age^13^, they do not exhibit an antiviral state. The cerebellum harbors diverse astrocyte and neuron subtypes, with Bergmann glia in contact with Purkinje Cells and interneurons, velate astrocytes with granule cells, fibrous astrocytes in the white matter, and astrocytes and neurons within the deep cerebellar nuclei (DCN)^35–37^. Cluster analysis of 23,344 cerebellar astrocytes combined with spatial transcriptomics analysis in adult mice^38^ identifies 2 Bergmann glia states (clusters 0&2) that differ in the expression level of neuroregulatory *Gpc5*^39,40^ and vasculature protective *Agpt1*^41^ factors. Velate and fibrous astrocytes in the granule cell layer and white matter (cluster 1), and astrocytes in the DCN (cluster 3) are also identified (Fig. 1F & S3E-H & Table S2). We find significant up-regulation of antiviral and reactive genes in all aged astrocyte sub-sets (Fig. 1G,H & S2H & Table S2), and Stat1 protein, a transcription factor that promotes antiviral gene expression^42^, is detected in Bergmann glia in the aged cerebellum, but not adult cerebellum or cortex at any age (Fig. 1I,J). Thus, antiviral signatures are not sub-set specific, but rather, a general characteristic of astrocytes in the aged cerebellum.

We next asked if regional astrocyte antiviral signatures are elicited progressively or whether they occur only in advanced age. We find enrichment for immune-related pathways and increased expression of 3 antiviral genes (*Ifit3, Ifit3b*, and *Casp1*) in cerebellar astrocytes compared to cortical already in adult (4mo) mice, using previous translatome data^13^ (Fig. S1G,H & Table S2). Further, the expression level of Interferon-related genes is higher in cerebellar astrocytes, compared to cortical, in adulthood (2 - 8.6 mo), but differences become more pronounced in advanced age (19.8 – 26.7mo), using astrocyte spatial transcriptomics data^43^ (Fig. 1K,L & Table S1). Inflammatory activity of Interferon Regulatory Factors (IRF) and NF-kB is also enriched in the cerebellum over the cortex, irrespective of age (Fig. 1M). Aged cerebellar astrocytes also have dysregulated expression of genes associated with Interferonopathies, whereas cortical do not, and broader changes in neurodegenerative disease genes (Fig. S4A,B, Table S2). In sum, the cerebellum as a compartment, and cerebellar astrocytes, have an immune-activated state throughout the lifespan that intensifies with age, that highlights a potential role for antiviral pathways in regulating aging changes in this brain region.

### Motor deficits in aging

The cerebellum plays a fundamental role in the coordination of limb movement, posture, and balance^8,9^, and aged human individuals experience motor deficits^5,7,44^. Analysis of data from the Rancho San Bernardo Healthy Aging Study^45^ demonstrates decreased limb strength, and slower, more imprecise performance in a narrow-line walking task in older individuals (Fig. 2A-C, S5A & Table S3). This prompted us to analyze motor function in adult and aged mice. Aged mice show decreased limb grip strength compared to adults (4mo), consistent with sarcopenia^1,46^ and with human data (Fig. 2A,D & S5A). Despite that, aged mice have increased body weight and equivalent basal locomotor activity to adults, and thus do not display sickness (Fig. S5B,C & Table S3). To assess motor function and locomotor strategy on a narrow walk task, mice are recorded while crossing elevated beams of decreasing width. All mice complete the task; however, we detect worse performance in aged mice, which cross 15-, and 10-mm-wide beams more slowly, and make more mistakes, with increased time stopped on the beam compared to adults (Fig. 2E-G). Differences are notable in the first trial on the 15-mm beam, and across trials with the 10-mm beam, suggesting aged mice are able to improve on a wider, easier beam. To investigate age-related adaptation in locomotor strategy, we performed kinematic analysis of mice crossing the 10-mm-wide beam and determined locomotion-related parameters. Principal Component Analysis finds differences in the strategy employed by aged mice; aged mice take slower, shorter steps with both paws (increased cycle, decreased displacement), and make different tail movements, with larger tail deviation and reduced tail velocity, perhaps to maintain balance^47^. Aged mice have hunched posture at the onset of each step, indicated by larger angle values between the base of the tail and the nose (Fig. 2H-K & S5D-I & Movie S1 & Table S3).

**Figure 2.**
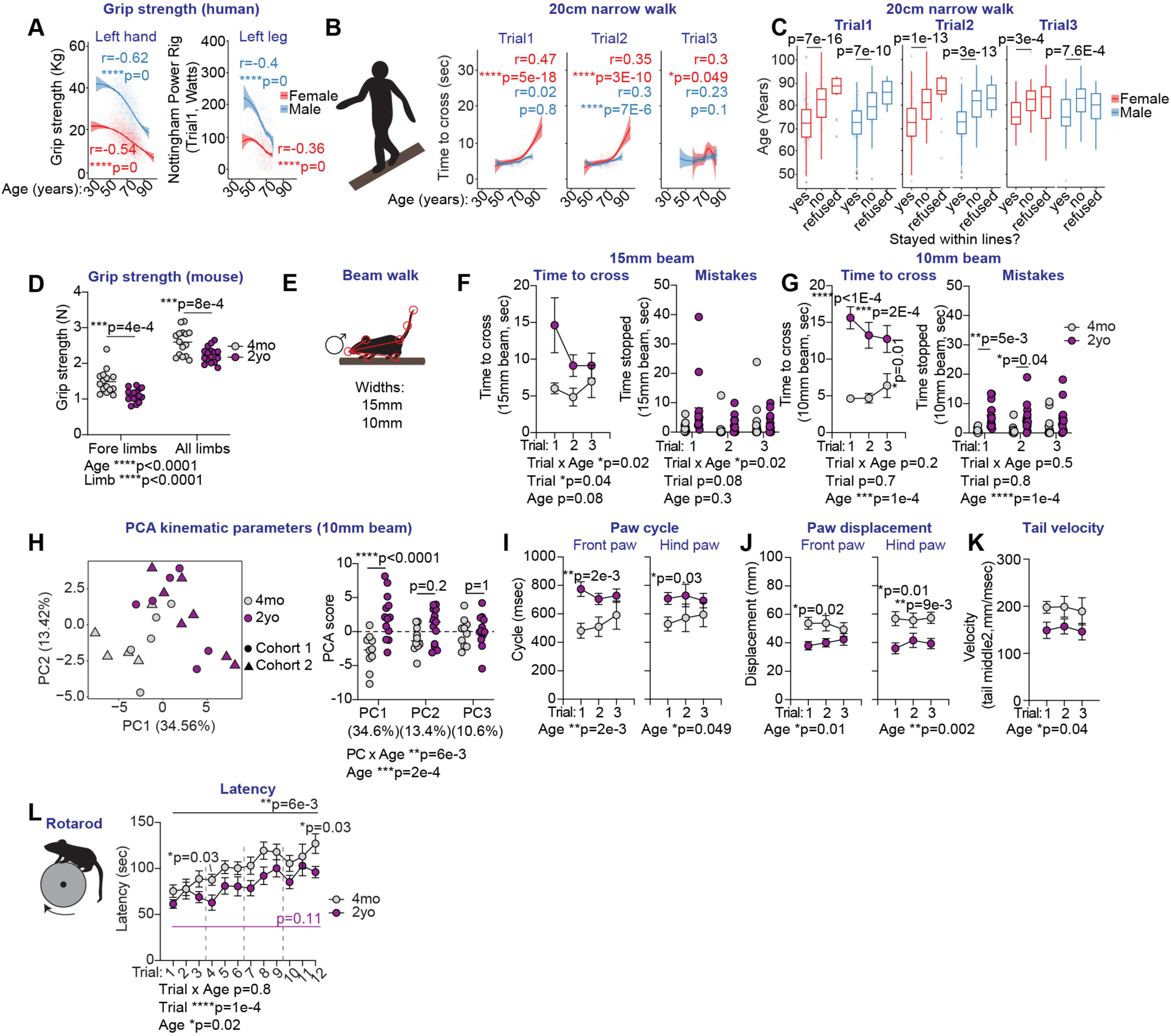
Motor deficits in aging. (A-C) Data are from the Rancho San Bernardo Study of Healthy Aging. (A,B) Correlation plots show individual data points and trend line indicating the relationship between grip strength of the left hand, and strength of the left leg with age (A), or between the time to cross a narrow line and age (B). (C) Box plots show the age distribution of human individuals that completed a narrow walk task without mistakes (‘Yes’), with mistakes (‘No’), or that did not perform the task (‘refused’). (D) Grip strength of fore limbs and all limbs in 4mo (adult) and 2yo (aged) WT male mice. Each circle corresponds to the average of 3 independent trials per mouse. (E-K) 4mo (adult) and 2yo (aged) WT male mice are analyzed while crossing 15mm, and 10mm-wide beams. (F,G) Average time to cross each beam, and total time stopped while crossing each beam is shown for every trial. Amount of time stopped indicates mistakes in this task. (E,H-K) Kinematic analysis of 4mo (adult) and 2yo (aged) WT male mice crossing the 10mm-wide beam. (H) Principal component analysis (PCA) using 45 kinematic parameters. PCA coordinate values for PC1, 2, and 3 are compared between mice of different ages. Percentages indicate the fraction of variability in the dataset explained by each component. (I) Paw cycle, (J) paw displacement, and (K) tail velocity of the tail middle 2 segment. (L) Latency of 4mo (adult) and 2yo (aged) WT male mice on the accelerating rotarod. (A-L) Statistical comparisons and kinematic data are provided in a Supplementary table. (A,B) Pearson correlation analysis, (C) Wilcoxon rank-sum test, (D,H) 2-way ANOVA Sidak’s multiple comparisons test, (F-G,I-L) RM ANOVA Sidak’s multiple comparisons test. (A) Number of data points: 7,509 (hand), 1,691 (leg), (B) 110-526 (time to cross), (C) 732 (task performance). (D) 15-16 mice/group. (F,L) 10-13 mice/group. (D,F,G) Horizontal bars indicate the mean, (F,G,I-L) mean ± S.E.M.

We also evaluated age-related motor adaptation with the accelerating rotarod, and find worse performance of aged mice compared to adults consistent with previous reports^48,49^. In addition, aged mice have reduced ability to improve on the rotarod, with significant improvement in adult mice, but not in aged, when comparing the first and last trial of this task (Fig. 2L & S5J & Table S3). Overall, we identify decreased muscle strength, adaptation in locomotor strategy and worse motor performance in aged mice that resembles that in cerebellar disease^50,51^ and in aged human individuals. This suggests involvement of the cerebellum, alongside other aging changes, in age-related motor dysfunction^44^.

### Type I IFN signaling promotes peripheral inflammation and motor deficits in aged mice

We next set out to test if blocking peripheral inflammation in aged mice can improve motor function and alleviate astrocyte aging changes. Given the up-regulation of antiviral signatures in aging, we focused on the type I Interferon (IFN) antiviral pathway. Aged mice were intraperitoneally (i.p.) treated with control or type I IFN receptor 1 (IFNAR1) blocking antibodies for 60 days, with IFNAR1 blockade sufficient to reduce hepatic expression of inflammation- and aging-related transcripts *Apol9b* and *Irf7*^16^ at the experimental endpoint, demonstrating effectiveness (Fig. 3A,B & S6A,B). To elucidate if peripheral blockade alleviates astrocyte phenotypes, we performed RT-qPCR analysis of cerebellar astrocyte nuclei (Fig.S2A-C & ^34^), finding equivalent antiviral gene expression in IFNAR1 and control groups (Fig. 3C & S6C). Reactivity-associated *Gfap*, and complement gene *C4b* are elicited in aged astrocytes^13,14^. We observe equivalent *C4b*, but significant reduction in *Gfap* transcripts in IFNAR1-treated mice compared to controls, although this does not correspond with reduced Gfap immunoreactivity (Fig. 3C & S6C-E). Thus, IFNAR1-blockade in aged mice alleviates inflammatory signaling in the periphery, and *Gfap* induction in astrocytes, but does not impact astrocyte antiviral signatures. Functionally, we find no treatment effect on basal locomotor activity or body weight (Fig. S6F,G & Table S3). However, IFNAR1-treated mice move faster than controls across the 15-mm-wide beam, although with similar accuracy (Fig. 3D-E & S6H,I & Movie S2). No differences are observed in the 10-mm-wide beam, which is more challenging at this age (Fig. S6I). In association with improved performance, we find changes in locomotor strategy; IFNAR1-treated mice take faster steps (decreased cycle), have increased tail velocity, and normalize their posture, compared to controls (Fig. 3F-H & S6J-M & Table S3). Thus, neutralization of peripheral IFNAR1 alleviates hepatic inflammation, and improves motor function in aged mice.

**Figure 3.**
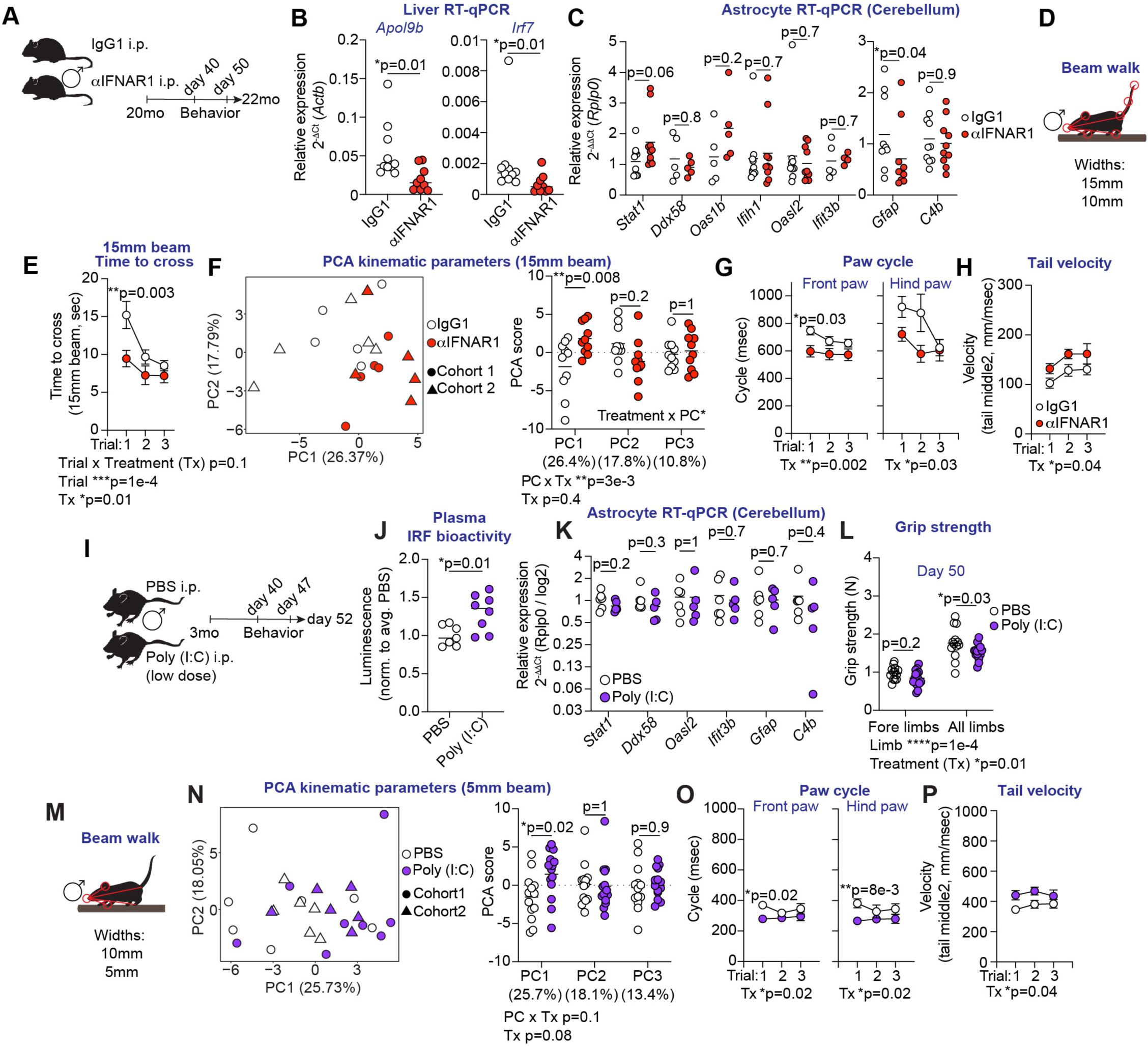
Type I IFN signaling promotes peripheral inflammation and motor deficits in aged mice. (A-H) 2yo (aged) WT male mice are intraperitoneally (i.p.) treated with IgG1 control or anti-IFNAR1 antibodies for 60 days. (B) Expression level of age-associated *Apol9b* and *Irf7* normalized to *Actb*, in the liver of IgG1 and anti-IFNAR1 treated mice at the experimental endpoint. (C) RT-qPCR analysis of astrocyte nuclei in the cerebellum at the experimental endpoint using *Rplp0* as reference gene. (D-H) Mice are analyzed while crossing 15mm, and 10mm-wide elevated beams. (E) Average time to cross the 15mm-wide beam for every trial. (F) PCA using 45 kinematic parameters obtained after analysis of the 15mm-wide beam. PC coordinates are compared between treatment groups. Percentages indicate variability explained by each component. (G) Per-step paw cycle, and (H) tail velocity of the tail middle 2 segment. (I-P) 3mo (adult) WT male mice are i.p. treated with PBS or a low dose of Poly (I:C) (0.4 mg/kg) for 52 days. (J) Interferon Regulatory Factor (IRF) bioactivity in plasma 3hrs after the last injection. (K) RT-qPCR analysis of astrocyte nuclei in the cerebellum at the experimental endpoint using *Rplp0* as reference gene. (L) Grip strength of fore limbs and all limbs in PBS or Poly (I:C) treated mice. Each circle represents the average of 3 independent trials per mouse. (M-P) Mice are analyzed while crossing 10mm, and 5mm-wide beams. (N) PCA using 45 kinematic parameters obtained after analysis of the 5mm-wide beam. PC coordinates are compared between treatment groups. Percentages indicate variability explained by each component. (O) Per-step paw cycle, and (P) tail velocity of the tail middle 2 segment. (B,C,J) Mann-Whitney tests, (E,G-H, O-P) RM 2-way ANOVA or (F,L,N) 2-way ANOVA with Sidak’s multiple comparisons test. (B,E,F-H) 10 mice/group, (C) 5-10 mice/group, (J) 7-8 mice/group, (K) 5-6 mice/group, (L-P) 13-14 mice/group). (B,C,F,J,K,L,N) Horizontal bars indicate the mean, (G,H,O,P) mean ± S.E.M.

We next asked if chronic viral inflammation at a low-level is sufficient to elicit motor deficits and astrocyte aging changes in adult mice. We focused on low-level inflammatory signaling to model what occurs in aging, and to avoid sickness. Adult (3mo) mice received repeated i.p. injections with PBS or with the viral pathogen-associated molecular pattern Poly (I:C) for 52 days, at a dose that is 10-30 times lower than common doses^52–54^ (Fig. 3I). Low-dose Poly (I:C) treatment induces IFN signaling in plasma, with increased IRF bioactivity, but not NF-kB, compared to controls (Fig. 3J & S7A). In the brain, we detect equivalent level of antiviral, *Gfap* and *C4b* transcripts in cerebellar astrocytes in both groups, and no differences in Gfap immunoreactivity across brain regions (Fig. 3K & S7B-D). Low-dose Poly (I:C) treatment does not induce sickness, with no impact on body weight nor basal locomotor activity (Fig. S7E-F), but it does lead to decreased grip strength of all limbs, similar to aged mice and elderly humans (Fig. 3L). Functionally, viral inflammation induces adaptation in locomotor strategy; Poly (I:C)-treated mice take faster steps (decreased cycle), and increase their tail movements, with increased tail velocity and tail deviation, compared to PBS controls, which points to a stimulatory effect, rather than detrimental (Fig. 3M-P & S7I-N & Table S3 & Movie S3). Locomotor adaptation induced after viral inflammation does not change the speed or accuracy when crossing elevated beams, and it does not affect performance or improvement in the rotarod task (Fig. S7H,O). Thus, low-level peripheral inflammation alone is insufficient to induce astrocyte antiviral signatures or motor deficits in adult mice.

Overall, we find that peripheral low-level inflammation does not elicit an antiviral state in astrocytes, but induces age-dependent motor adaptation with a stimulatory effect in adults, and detrimental impact in advanced age. This suggests that peripheral inflammation alone is not causative, but may act in synergy with other aging changes to influence motor function.

### Astrocyte Stat1 promotes motor deficits during chronic inflammation

Stat1 is an antiviral transcription factor that is up-regulated in cerebellar astrocytes during advanced age (Fig. 1I). To elucidate if astrocyte Stat1 is sufficient to promote motor deficits and astrocyte aging changes in adulthood, juvenile mice were retro-orbitally (r.o.) injected with adeno-associated viruses (AAVs) that cross the blood-brain barrier^55^, and that encode HA-tagged Stat1 (Stat1-HA) or a control protein (spaghetti monster fluorescent protein, smFP^56^) under an astrocyte promoter^57^. At 4 months of age, ∼60 % of astrocytes in the cerebellum express either protein, with very low off-target cells (Fig. 4A-D). RT-qPCR of cerebellar astrocyte nuclei detects 3-16-fold increase in *Stat1* transcripts in Stat1-HA compared to smFP mice, but no induction of antiviral transcripts, and reduced *Gfap*, perhaps due to competition with Stat3^42,58^ (Fig. 4D & S8A). We do not observe differences in body weight, basal activity, motor performance or locomotor strategy on the narrow beam between both groups (Fig. S8B-J & Movie S4). Similarly, we do not find differences in rotarod performance, although significant improvement between the first and last trial of this task is observed in smFP controls, but not in Stat1-HA (Fig. 4G & S8K & Table S3). Thus, overexpression of Stat1 in astrocytes does not substantially recapitulate aging changes in adult mice in homeostasis.

**Figure 4.**
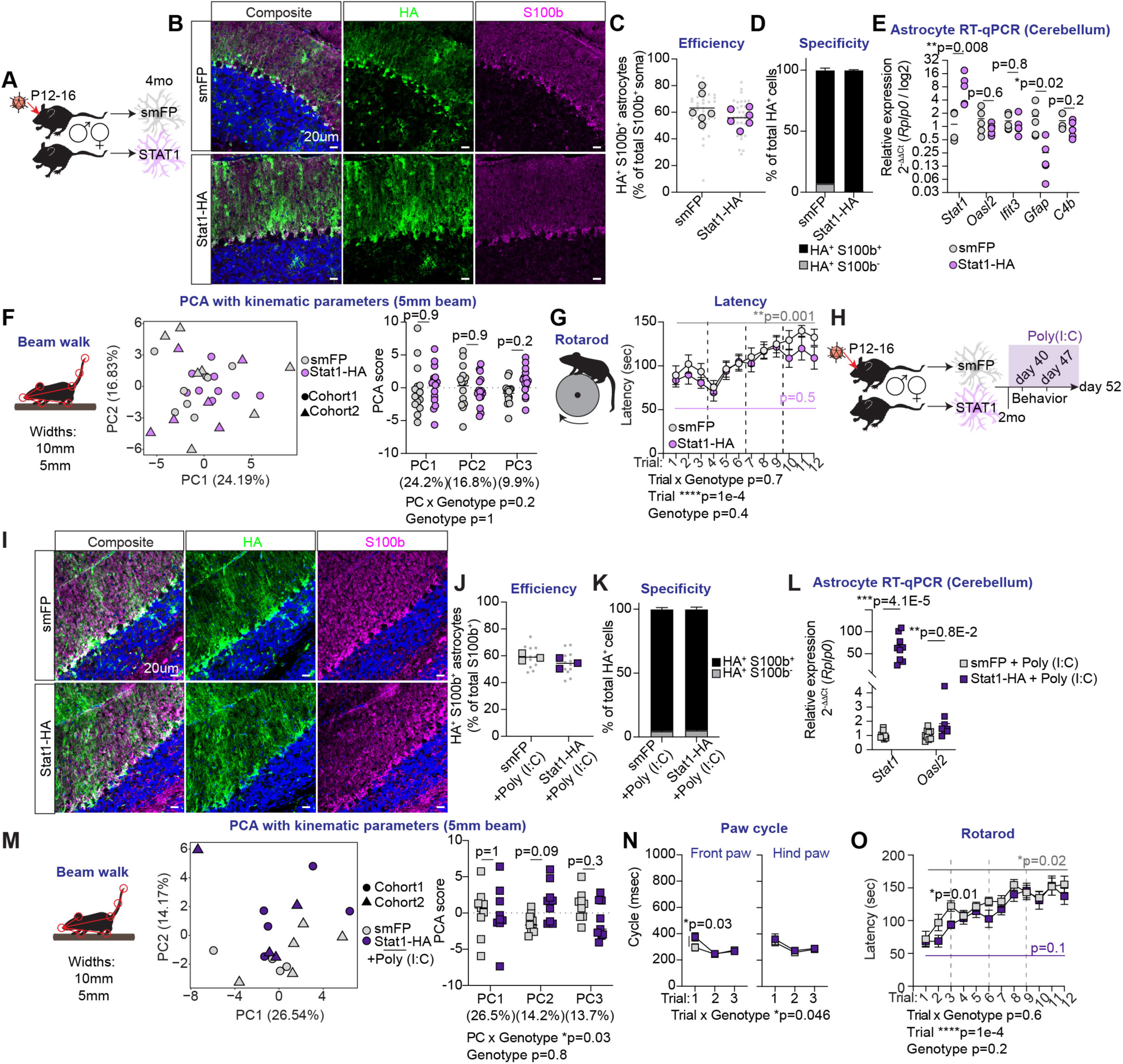
Astrocyte Stat1 promotes motor deficits during chronic inflammation. (A-G) Male and female WT mice are retro-orbitally (r.o.) injected AAVs at post-natal days 12-16. Behavioral testing is performed at 4mo, and the experiment terminated at 4.5mo. (B-D) Staining with HA antibody detects smFP and Stat1-HA co-localization with S100b in the cerebellum at 4.5mo. (C) Percentage of S100b+ soma that are HA+ (efficiency). Small grey dots are individual images; grey and pink circles are mouse averages. (D) Percentage of HA+ soma that are S100b+ or S100b- (specificity). (E) RT-qPCR analysis of astrocyte nuclei in the cerebellum at the experimental endpoint using *Rplp0* as reference gene. (F) Mice are analyzed while crossing 10mm, and 5mm-wide beams. PCA using 45 kinematic parameters after analysis of the 5mm-wide beam. PC coordinates are compared between genotypes. Percentages indicate variability explained by each component. (G) Latency on the accelerating rotarod. (H-O) Male and female WT mice are r.o. injected AAVs at P12-16. Starting at 2mo, both groups of mice are i.p. treated with low dose Poly (I:C) (0.4 mg/kg) for 52 days. Behavioral testing is performed between day 40 and 47 of treatment, and the experiment terminated at 4.5mo. (I-K) HA staining detects smFP and Stat1-HA co-localization with S100b in the cerebellum at 4.5mo. (J) Percentage of S100b+ soma that are HA+. Small grey dots are individual images; grey and violet squares are mouse averages. (K) Percentage of HA+ soma that are S100b+ or S100b-. (L) RT-qPCR analysis of astrocyte nuclei in the cerebellum at the experimental endpoint using *Rplp0* as reference gene. (M,N) Mice are analyzed while crossing 10mm, and 5mm-wide beams. (M) PCA using 45 kinematic parameters obtained after analysis of the 5mm-wide beam. PC coordinates are compared between genotypes. Percentages indicate variability explained by each component. (N) Per-step paw cycle. (O) Latency on the accelerating rotarod. (E, L) Mann-Whitney tests, (F,M) 2-way ANOVA or (G,N,O) RM 2-way ANOVA with Sidak’s multiple comparisons test. (C,D) 6 mice/group, (E) 5 mice/group, (F,G) 14-15 mice/group, (J,K) 3 mice/group, (L-O) 9 mice/group. (C,E,F,J,L,M) Horizontal bars indicate the mean, (D,G,K,N,O) mean ± S.E.M.

Since astrocyte changes occur in parallel with peripheral inflammation, we reasoned that Stat1 in astrocytes may exert its functions during chronic inflammation. To test the functional role of Stat1 in this context, we overexpressed smFP or Stat1-HA in astrocytes in juvenile mice, as before, and, starting at 2mo, both groups of mice were repeatedly treated with low-dose Poly (I:C) for 52 days (Fig. 4H). At endpoint, ∼58 % of cerebellar astrocytes overexpress either protein, with high cell-type specificity (Fig. 4I-K). RT-qPCR analysis of cerebellar astrocyte nuclei finds 20-128-fold induction of *Stat1* transcripts in Stat1-HA compared to smFP, as well as increased *Oasl2,* which is reduced in cerebellar astrocytes in Stat1^-/-^ mice (Fig. 4L & S9A-D). Other aging-associated genes are not consistently changed after Stat1 overexpression (Fig. 4J & S9A-C). Thus, peripheral inflammation amplifies Stat1 expression, from 3-16-fold to 20-128-fold increase over smFP control (Fig. 4E,L), along with *Oasl2* level following Stat1 overexpression. The limited effect on additional antiviral genes likely reflects the absence of IFN signaling in astrocytes after low-level peripheral inflammation (Fig. 3K). We find no differences in body weight, grip strength, basal activity, nor in performance or accuracy while crossing narrow beams between both groups (Fig. S9E-H). However, Stat1 overexpression in combination with Poly (I:C) induces changes in locomotor strategy, including slower fore paw steps (increased cycle), like observed in aged mice, and increased tail movements (tail distance) (Fig. 4M,N & S9I-M & Movie S5 & Table S3). In addition, Stat1-HA mice with Poly (I:C) perform worse than smFP controls on the rotarod, particularly in early trials, and have reduced improvement between the first and last trials of this task (Fig. 4O & S9N & Table S3). Overall, we identify a functional role for astrocyte Stat1 in the promotion of motor deficits in crosstalk with peripheral inflammation.

### Systemic high-level viral inflammation does not drive regional astrocyte antiviral state

Given the functional relevance of type I IFN signaling and Stat1 in aging (Figs. 3&4), we next set out to elucidate the triggers of astrocyte antiviral state in the cerebellum. We first asked if astrocytes in the cerebellum respond to systemic viral challenge and elicit aging changes. Adult (3mo) and aged (20mo) mice were i.p. infected with Lymphocytic choriomeningitis virus (LCMV), which drives high-level inflammation engaging innate and adaptive immune pathways^59,60^. RT-qPCR analysis of cerebellar astrocyte nuclei finds significant impact of infection on *Stat1* and *C4b* level particularly in aged mice, and a trend in adults, at day 7 post-infection. *Gfap* mRNA level or immunoreactivity is unchanged by infection at this time-point (Fig. S10A-E). Thus, cerebellar astrocytes upregulate aging-associated genes in response to a recent viral infection, particularly in aging, when signaling is basally higher. We next wondered if cerebellar astrocytes are more responsive to high-level viral inflammation than cortical. Viral infections induce copious type I IFNs early after infection^59^, thus, to emulate early stages of infection, adult (2.5mo) mice were i.p. injected with PBS or with Poly (I:C) for 3 consecutive days at a dose 10-times higher than used in our low-level inflammation protocol (Fig. S10F). High-dose Poly (I:C) increases plasma IRF bioactivity in higher magnitude than low-dose compared to controls, and it does not elicit NF-kB. It also induces body weight loss, indicative of sickness, which is not observed with the low dose or in aged mice (Fig. S10G,H). High-level inflammation induces *Ifnb1* transcripts in the choroid plexus^61^, and Stat1 immunoreactivity across brain regions (Fig. S10I-K,M). In astrocytes, Poly (I:C) treatment increases the expression of antiviral genes and *C4b* both in cortical and cerebellar astrocytes, while Gfap immunoreactivity remains unchanged compared to PBS controls (Fig. S10L-O). Thus, we find that the capacity to respond to high-level viral inflammation is not a unique trait of cerebellar astrocytes, which points to a role for local signaling in regional regulation.

### Local type I IFN signaling drives astrocyte regionality in aging

We next sought to elucidate if type I IFN signaling is required for cerebellar regionality, and whether in aging, the source of IFNs is localized within the cerebellum. We find decreased *Stat1* mRNAs in the cerebellum of adult (7-8mo) mice that lack functional type I IFN receptor (IFNAR1^-/-^), compared to WT controls. This reduction is not detected in the cortex, and is more pronounced in the Purkinje Cell layer of the cerebellum, which contains Bergmann glia and Purkinje Cell soma (Fig. 5A,B). To elucidate if local type I IFNs are drivers, aged mice were locally infused with control or anti-IFNAR1 antibodies into the cerebellum. Short-term IFNAR1-blockade reduces the level of *Stat1* mRNAs in the molecular and Purkinje cell layers, and alleviates the induction of multiple antiviral gene transcripts in astrocytes in the aged cerebellum. *Gfap* and *C4b* transcript abundances are unchanged by anti-IFNAR1 treatment (Fig. 5C-F & S11A). Thus, type I IFN signaling in the cerebellum sustains antiviral gene expression in astrocytes, but is not responsible for all aging changes.

**Figure 5.**
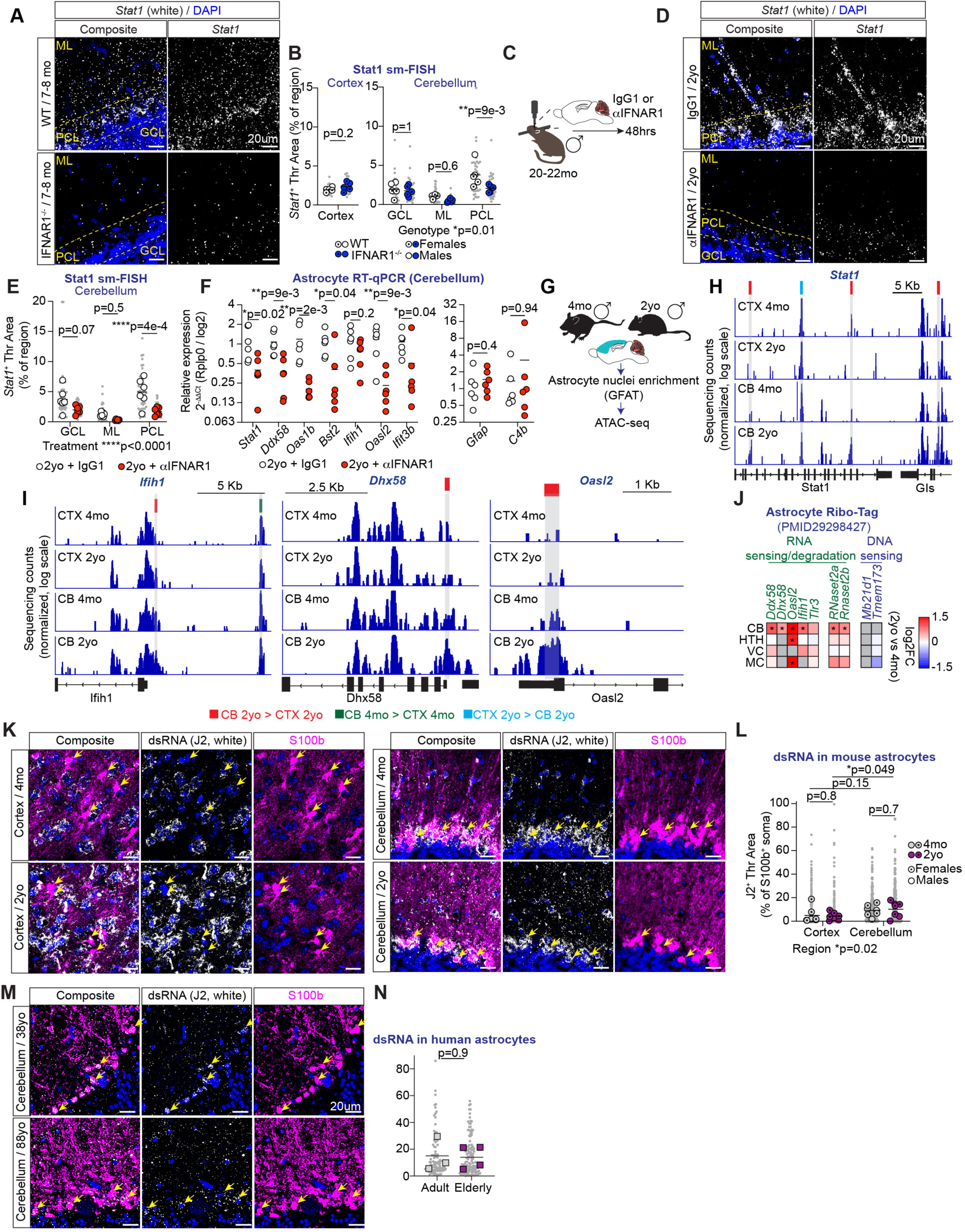
Local type I IFN signaling and chromatin de-repression promote astrocyte antiviral state in the aged cerebellum. (A,B) Stat1 mRNAs detected using single-molecule Fluorescent In Situ Hybridization (sm-FISH) in the cortex and cerebellum of 7-8mo WT and Ifnar1-/- mice. Dotted lines indicate the cerebellar granule cell layer (GCL), molecular layer (ML), and Purkinje Cell layer (PCL). (B) Quantification of Stat1 signal in the cortex, and cerebellar layers. Grey dots are individual images; white and blue circles are mouse averages. (C-F) 20-22mo aged WT male mice are injected with IgG1 or anti-IFNAR1 into the cerebellum, and cerebella analyzed 48hrs after injection. (D) Stat1 mRNAs detected with sm-FISH in the cerebellum of IgG1 and anti-IFNAR1 injected mice. Dotted lines indicate GCL, ML, and PCL. (E) Quantification of Stat1 signal in cerebellar layers. Grey dots are individual images; white and red circles are mouse averages. (F) RT-qPCR analysis of astrocyte nuclei in the cerebellum at the experimental endpoint using *Rplp0* as reference gene. (G-I) Astrocyte nuclei in the cortex and cerebellum of 4mo and 2yo male WT mice (3 mice/group) are enriched using Glyoxal Fixed Astrocyte Transcriptomics followed by Assay for Transposase-Accessible Chromatin using sequencing (ATAC-seq). (H,I) Representative DNA library tracks with differentially accessible regulatory regions between indicated comparisons. For each gene, all tracks use the same scale, allowing visual comparison. (J) Heatmaps show log2 fold changes in gene expression of genes involved in RNA sensing and degradation, and DNA sensing in astrocytes in the cerebellum (CB), hypothalamus (HTH), visual cortex (VC), and motor cortex (MC) using published Ribo-Tag data. (K-N) Detection of double stranded (ds)RNAs with the J2 antibody, and S100b protein with immunohistochemistry in the cortex and cerebellum of 4mo (adult) and 2yo (aged) WT male and female mice (K,L) and in human cerebellum (M,N). Arrows point at S100b+ astrocyte soma. (L,N) Quantification of J2 signal in S100b+ soma in K and M. Grey small dots are individual S100b+ soma; grey and magenta circles or squares are mouse or human averages. (M,N) Specimens are 34, 38, 41 years old (adults), and 80, 81, 85, 88 (elderly). (B-cortex, F,N) Mann-Whitney tests, (B-cerebellum, E) 2-way ANOVA with Sidak’s multiple comparisons test. (H-I) Differentially accessible regulatory regions (DARs) defined by adjusted p.val<0.05 and absolute fold change > 2. (A,B) 5 mice/group, (C-F) 6 mice/group, (G) 3 mice/group, (L) 7 mice/group, (N) 3-4 human individuals per group. (B,E,F,L,N) Horizontal bars indicate the mean.

We next hypothesized that microglia are a likely source, as they produce type I IFNs^24^, and promote immune gene expression in astrocytes^26,62^. To test this, microglia were depleted from aged mice using chow containing the CSF1R antagonist PLX3397 (PLX) for 56 days^63^ (Fig. S11B-F). RNA-seq analysis of cerebellar astrocyte nuclei finds PLX-dependent gene expression shifts, but does not reverse most aging-changes, including antiviral and reactive genes (Fig. S11G-K & Table S2). Thus, microglia are not sufficient to sustain antiviral or reactive signatures in aged cerebellar astrocytes. Overall, we find that type I IFN signaling drives astrocyte regionality, and that, local signaling promotes exacerbated antiviral signatures in aging.

### Chromatin de-repression of antiviral gene regulators amplifies regionality in aging

We next focused on astrocyte-intrinsic changes that could explain aging signatures. We asked if antiviral genes are epigenetically de-repressed in cerebellar astrocytes relative to cortical, leading to increased basal expression and in aging. To define the chromatin landscape of regional astrocytes, we performed Assay for Transposase-Accessible Chromatin using sequencing (ATAC-seq)^64^ in astrocyte nuclei in the adult (4mo) and aged (2yo) cerebellum obtained using GFAT^34^ (Fig. 5G). ATAC-seq identifies regional differences that are more pronounced in advanced age, with small impact of age in chromatin accessibility within each region (Fig. S12A-G & Table S4). We find increased accessibility at regulatory regions of multiple antiviral genes including *Irf1*, *Irf7*, *Ifnar1*, *Ifngr1,* and *Ifnk* in cerebellar astrocytes, compared to cortex, in 4mo and 2yo mice (Fig. S12H-J). Some regions are only de-repressed in advanced age, including those regulating *Stat1*, genes encoding cytosolic double-stranded (ds)RNA sensors (*Ifih1*, *Dhx58*, and *Oasl2)*, and *Casp1*, which are also increased at the transcript or protein level (Fig. 5H,I). Intriguingly, *Ifih1* transcripts are not reduced after IFNAR1 blockade (Fig. 5F & S11A), suggesting that increased chromatin accessibility can drive initial expression of regional genes, prior to amplification by IFN-I signaling. Thus, we find increased de-repression of antiviral gene regulatory regions in cerebellar astrocytes compared to cortical during adulthood, that is exacerbated in aging. Alongside dsRNA sensors, increased expression of ribonucleases *RNaset2a*, and *RNAset2b* is also detected (Fig. 5J & S11L). We also find dsRNA accumulation in cerebellar astrocytes, in adulthood and aging, but not cortical (Fig. 5K,L & S11M,N), suggestive of increased responsiveness against self-dsRNAs by aged cerebellar astrocytes. DsRNAs are also detected in human Bergmann glia in the adult and aged cerebellum (Fig. 5M,N & S11O). DNA sensors cGAS and STING (*Mb21d1* and *Tmem173*) are not detected in mouse cerebellar astrocytes (Fig. 5J & S11L)^26^. In sum, heightened antiviral signaling in aged cerebellar astrocytes can be explained by increased chromatin accessibility of transcription factor Stat1, and heightened capacity to sense and degrade self dsRNAs, which, together, can amplify local IFN-I signaling.

Our studies show that a classical antiviral pathway (IFN-I) and the antiviral gene Stat1 in astrocytes promote motor decline similar to motor deficits observed in aged human individuals. We establish an immune activated state in astrocytes in the cerebellum, but not in cortex, that arises in adulthood and that is exacerbated in aging. We discover that such state is a result of persistent type I IFN signaling within this brain region both in adulthood and in aging, which may drive unique conformation of the chromatin landscape, making cerebellar astrocytes more permissive to sensing and amplification of inflammation. In aging, astrocyte regional differences are accentuated by de-repression of additional regulators of the immune response, including transcription factor Stat1, and dsRNA sensing machinery that, together, can amplify antiviral gene programs^26,65,66^. We establish that immune-astrocyte communication is functional in brain aging and functional decline, and that targeting the type I IFN pathway can improve motor health in advanced age.

## Acknowledgements

This work is supported by R21 NS137659, Chan Zuckerberg Initiative, and Roger Guillemin Chair to NJA, NOMIS NeuroImmune Initiative to NJA, DCH, and SMK. LL-B is supported by Kavli Innovative Research Grant, Salk Women & Science Award, AHA BHCI Collaborative Grant, George E Hewitt Postdoctoral Fellowship, and NIH-NlA San Diego Nathan Shock Center P30 AG068635. ART is supported by Salk Alumni Fellowship Award, and Salk Women & Science Research Award. BYC is supported by NIH grant 5T32GM133351. EA is supported by NIH grants R01 NS128898, U19 NS112959, R01 NS111479, and The Callahan Foundation. This work is supported by Animal Resources Department, In Vivo Scientific Services (IVSS), and The Razavi Newman Integrative Genomics and Bioinformatics Core (IGC) at the Salk Institute. This work is supported by the Flow Cytometry Core Facility of the Salk Institute (RRID:SCR 014839) with funding from NIH-NCI CCSG: P30 CA01495, and Shared Instrumentation Grants S10-OD023689 (Aria Fusion cell sorter) and S10 OD034268 (Thermo Fisher Bigfoot). This work is supported by the Gene Transfer, Targeting and Therapeutics Viral Vector Core (GT3) of the Salk Institute with funding from NIH-NCI CCSG: P30 CA01495, an NINDS R24 Core Grant and funding from NEI (R24NS092943). This work is supported by the Waitt Advanced Biophotonics Core Facility of the Salk Institute (RRID:SCR_014838) with funding from NIH-NCI CCSG P30 CA014195, NIH-NIA San Diego Nathan Shock Center P30 AG068635, The Henry L. Guenther Foundation and the Waitt Foundation. This publication includes data generated at the UC San Diego IGM Genomics Center utilizing an Illumina NovaSeq 6000, and an Illumina NovaSeq X Plus that were purchased with funding from a National Institutes of Health SIG grant (#S10 OD026929). Human tissue was received from the NIH NeuroBioBanks at the University of Miami, the Sepulveda Research Corporation, and the Harvard Brain Tissue Resource Center. This work is supported by the UC San Diego Biorepository and Tissue Technology Shared Resources (BTTSR) at the Moores Cancer Center. We thank all members of the Allen lab for providing feedback on this study, Dr John Teijaro at The Scripps Research Institute for kindly sharing Ifnar1^-/-^ and Stat1^-/-^ mice.

---

**Supplementary Figure 1 (Related to Figure 1).**
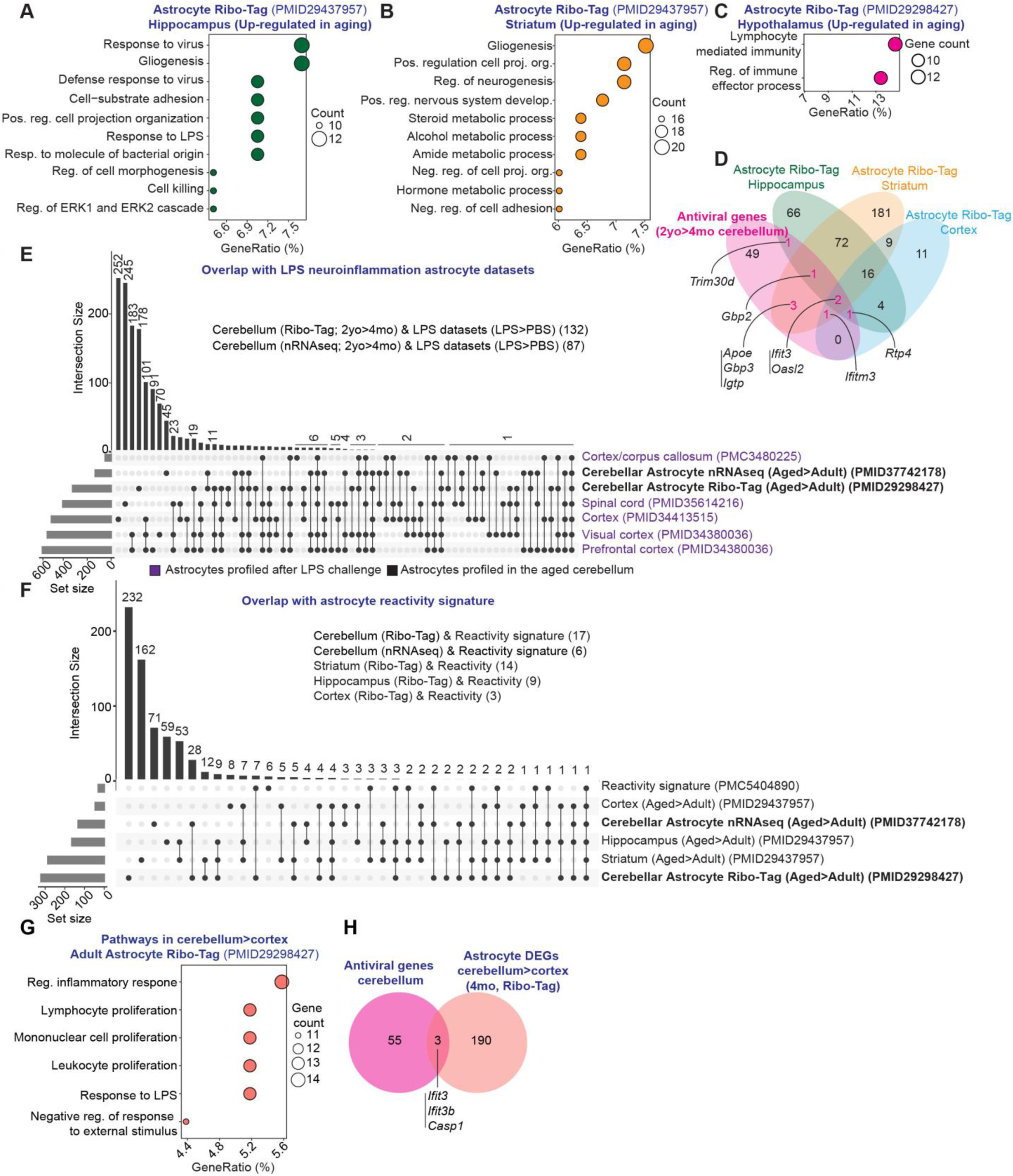
Astrocytes in the adult and aged cerebellum exhibit immune gene signatures. (A-C) Gene ontology (GO) pathway analysis using 2 published astrocyte Ribo-Tag RNA-seq datasets that profile astrocytes in the adult (2-4mo) and aged (2yo) hippocampus, striatum, and hypothalamus of WT male mice. DEGs up-regulated in 2yo vs 4mo were used as provided in the original study for hippocampus, and striatum. For hypothalamus an additional cut-off for absolute fold-change (FC)>1.5 was used to DEGs reported in the original study. Pathways with 10 or more DEGs overlapping with GO pathways is shown. GeneRatio refers to the percentage of DEGs, among all DEGs, that overlap with a GO pathway. (D) Venn diagram overlaps 58 antiviral genes up-regulated in aged cerebellar astrocytes with DEGs up-regulated in 2yo vs 2mo mice in astrocytes in the hippocampus, striatum, and cortex defined in PMID20437957. (E,F) UpSet plots overlap DEGs up-regulated in 2yo vs 4mo cerebellar astrocytes (Ribo-Tag and nRNA-seq datasets) with genes up-regulated in astrocytes after LPS-induced neuroinflammation (E), and with reactivity-associated genes (F). (G) GO pathway analysis using DEGs up-regulated in astrocytes in the cerebellum vs cortex in adult (4mo) mice. DEGs were previously identified with Ribo-Tag RNA-seq, and we included an additional cut-off for |FC|>1.5. (H) Venn diagram overlaps DEGs up-regulated in cerebellar vs cortical astrocytes in adult (4mo) mice with 58 antiviral genes up-regulated in aged cerebellar astrocytes.

**Supplementary Figure 2 (Related to Figure 1).**
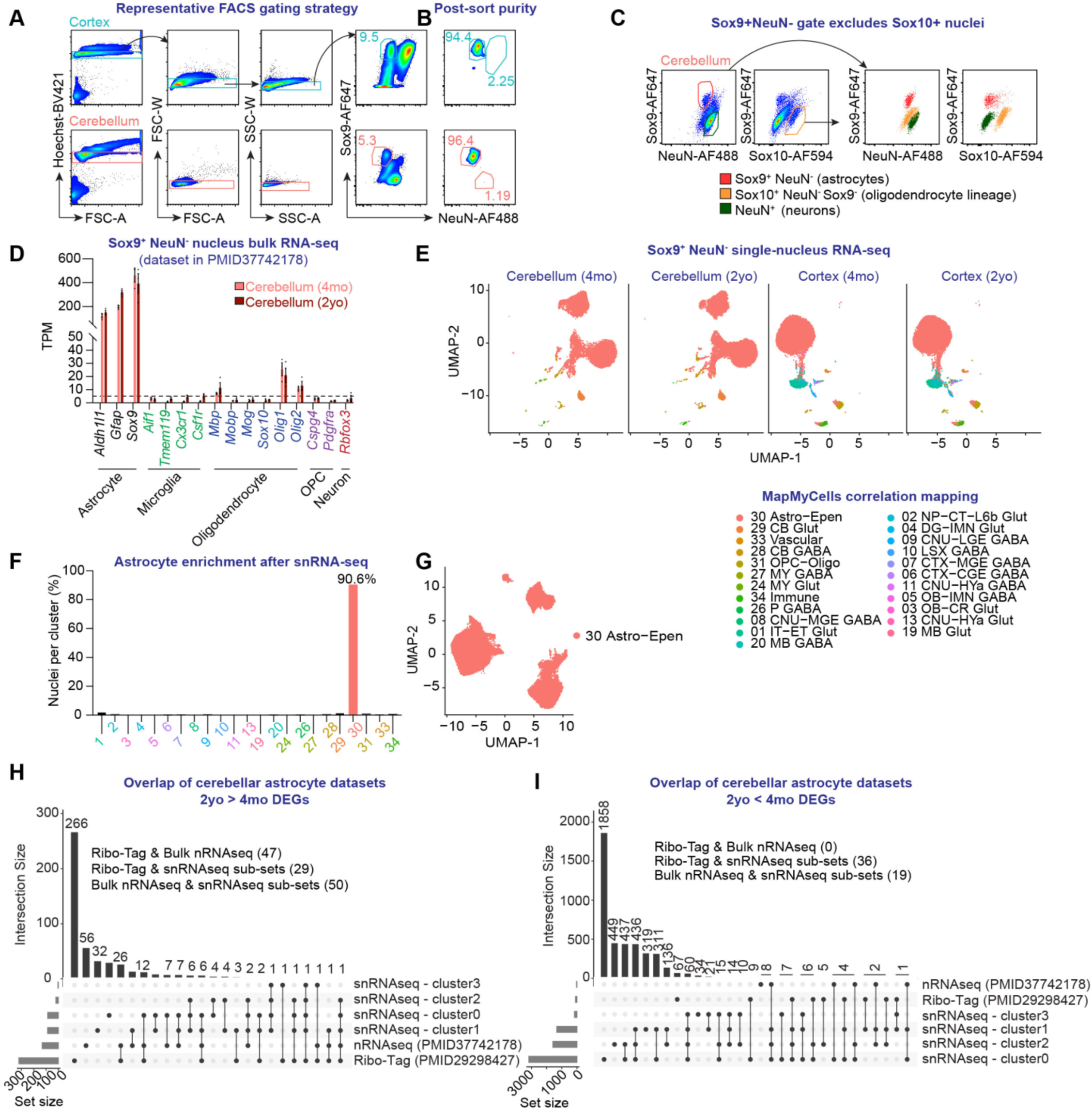
Enrichment of astrocyte nuclei using Glyoxal Fixed Astrocyte Transcriptomics. (A-G) Cortex and cerebellum are dissected from adult and aged mice and briefly fixed in glyoxal solution. Nuclei suspensions are obtained and stained with Hoechst, anti-NeuN, anti-Sox9 primary, and fluorophore-conjugated secondary antibodies. (A) Representative gating strategy to FACS-purify Sox9+NeuN- astrocyte nuclei. Rectangular gates are placed around single nuclei using FSC-A vs. Hoechst, FSC-A vs FSC-W and SSC-A vs SSC-W. A polygon gate is placed around Sox9+ NeuN- to gate on astrocyte nuclei. Gate numbers represent the percentage of single nuclei that are Sox9+ NeuN-. (B) Post-sort purity is confirmed after running a small volume of sorted sample through the FACS. Gate numbers are the percentage of sorted nuclei that are Sox9+NeuN-. (C) Sox9+NeuN- gating excludes Sox10+ oligodendrocyte, and NeuN+ neuronal nuclei. (D) We previously performed RNA-seq on Sox9+NeuN- FACS-purified nuclei in the cerebellum of 4mo (adult) and 2yo (aged) male mice. An expression cut-off of 5 transcripts per million (TPM) is used, which enriches astrocyte transcripts over those of microglia, oligodendrocyte, oligodendrocyte precursor cells (OPC), and neurons. (E-G) Single-nucleus (sn)RNA-seq is performed using Sox9+NeuN- FACS-purified nuclei in the cerebellum and cortex of 4mo (adult) and 2yo (aged) male mice. 2 mice per group are pooled into one biological sample. MapMyCells Correlation clustering (Allen Institute for Brain Science) detects a low number of neuron, microglia and oligodendrocyte nuclei (E,F) that are removed for downstream astrocyte analysis (G). (H,I) Upset plots compare all astrocyte cerebellar datasets used in this study. The degree of overlap among DEGs up-regulated (H) or down-regulated (I) in 2yo (aged) vs 4mo (adult) across datasets is shown. (D) Mean ± S.E.M.

**Supplementary Figure 3 (Related to Figure 1).**
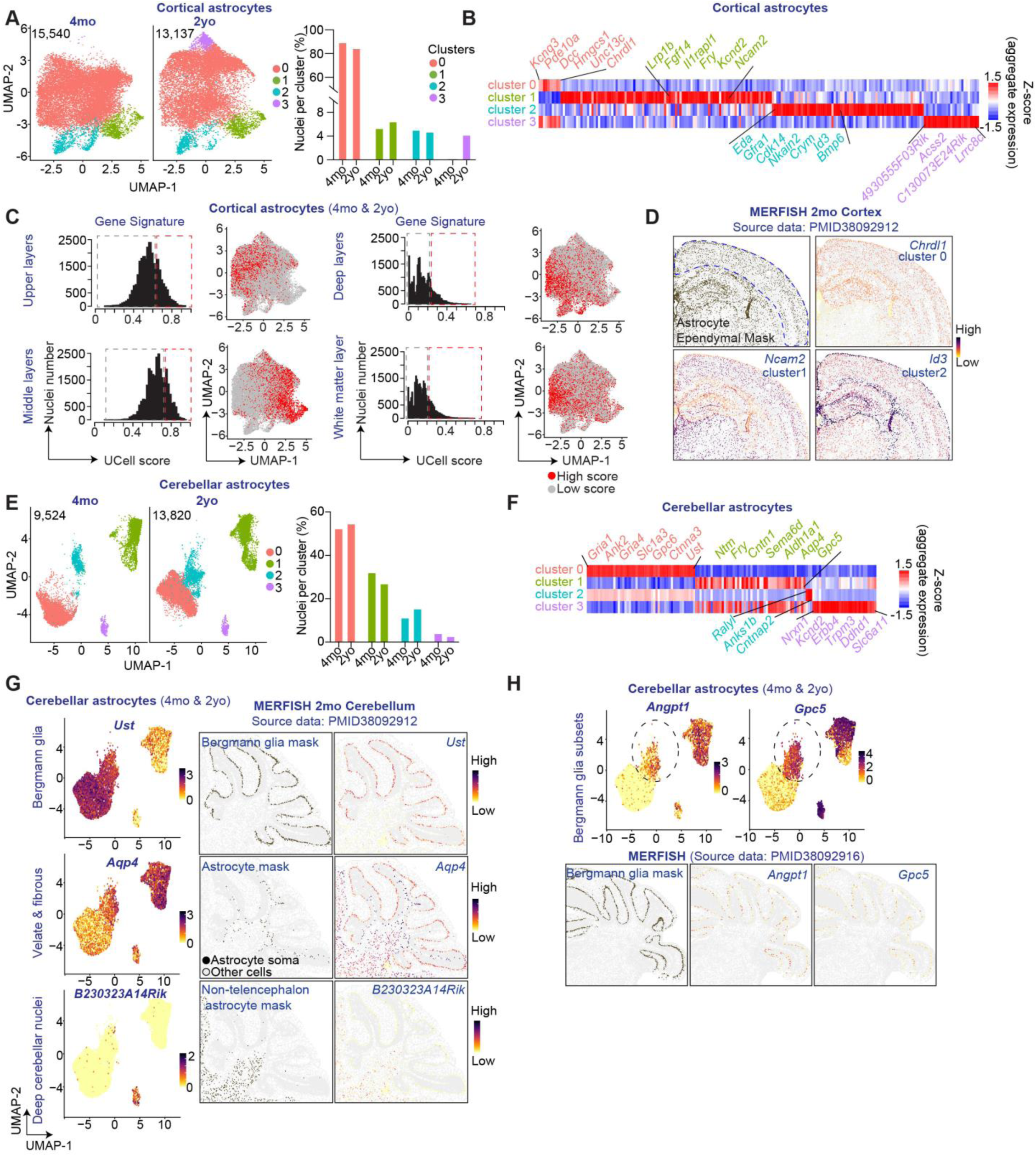
Astrocyte snRNAseq identifies sub-sets in the cortex and cerebellum. (A-H) Astrocyte snRNA-seq is performed using Sox9+NeuN- FACS-purified nuclei in the cerebellum and cortex of 4mo (adult) and 2yo (aged) male mice. 2 mice per group are pooled into one biological sample. (A) UMAP plots show 3 astrocyte clusters in the cortex of 4mo, and 4 in 2yo mice. Numbers in the upper left corner indicate the number of astrocyte nuclei analyzed. Bar graph shows nuclei proportions in each cluster and age. (B) Heatmap shows transcriptional markers of cortical subsets identified in (A). (C) Histograms show the distribution of UCell scores in cortical astrocytes for layer-enriched gene signature previously defined. Red rectangles mark astrocytes with 25% highest scores. UMAP plots show astrocytes in adult and aged mice. Astrocyte nuclei in red have highest UCell scores for each signature. (D) Representative images show mRNA level of cluster-defining genes *Chrdl1*, *Ncam2*, and *Id3* in the cortex of an adult (2mo) mouse. (E) UMAP plot shows 4 astrocyte clusters in the cerebellum of 4mo, and in 2yo mice. Numbers in the upper left corner indicate the number of astrocyte nuclei analyzed. Bar graph shows the proportion of nuclei in each cluster and age. (F) Heatmap shows transcriptional markers of cerebellar subsets identified in (E). (G,H) UMAP plots show relative expression of *Ust* in Bergmann glia, *Aqp4* in velate & fibrous astrocytes, and *B230323A14Rik* in astrocytes in the deep cerebellar nuclei (G). (H) Bergmann glia sub-sets can be distinguished with expression of *Angpt1* and *Gpc5*. Representative images show mRNA expression of subset-defining genes, and indicate localization of Bergmann glia in the Purkinje Cell layer, velate & fibrous in the granule cell layer and white matter, and in the deep cerebellar nuclei.

**Supplementary Figure 4 (Related to Figure 1).**
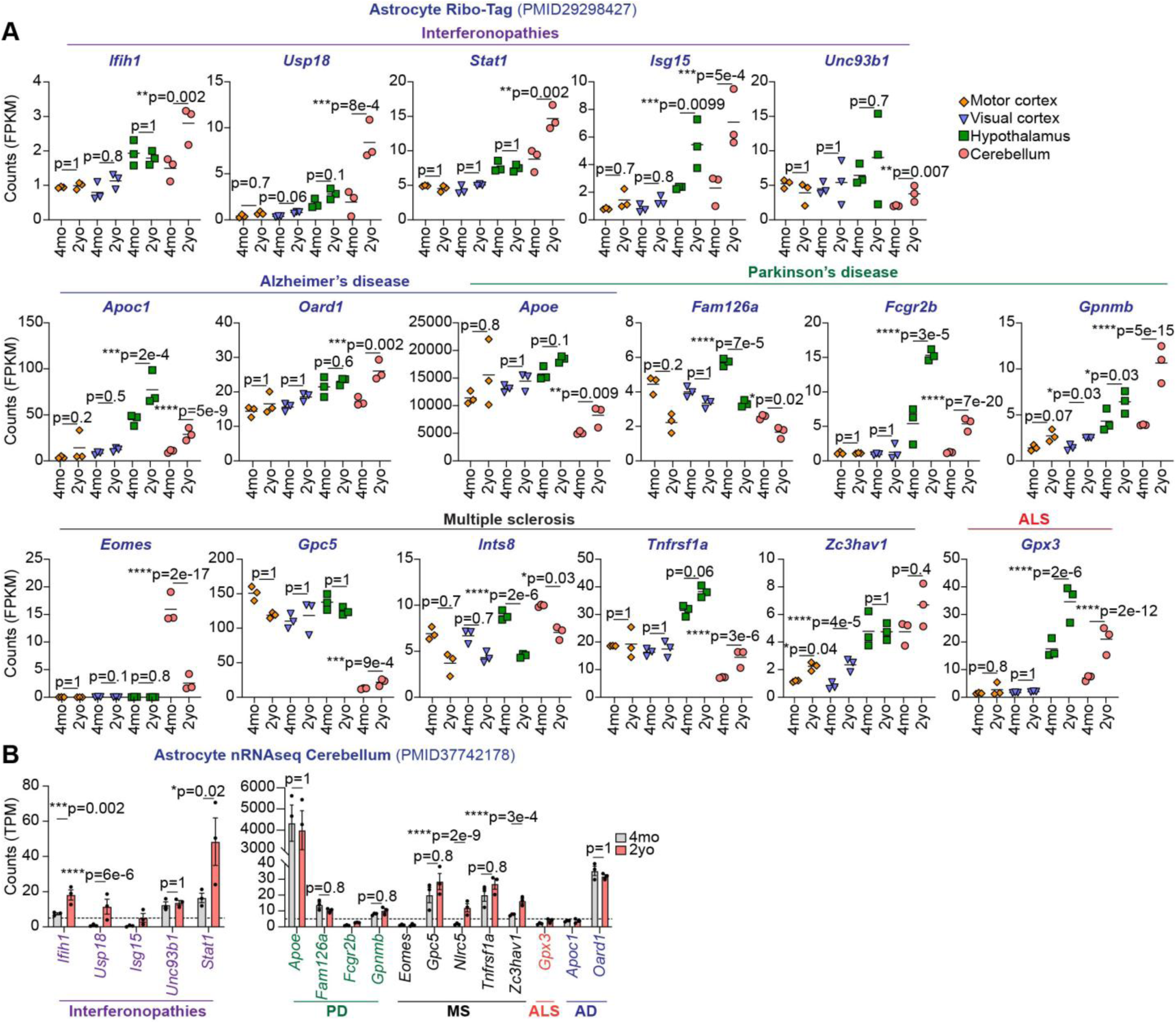
Expression of disease-associated genes in astrocytes in adulthood and in aging. (A) Expression level of genes associated with human interferonopathies and neurodegenerative diseases in astrocytes in the motor and visual cortex, hypothalamus and cerebellum of 4mo (adult) and 2yo (aged) WT male mice. Source data is astrocyte Ribo-Tag RNA-seq, and counts are in Fragments per Kilobase Million (FPKM). (B) Expression level of disease-associated genes in the cerebellum of 4mo (adult) and 2yo (aged) WT male mice. Source data is astrocyte nRNA-seq, and counts are in Transcripts per Million (TPM). (A,B) Adjusted p-values provided in the original publication (A), and obtained using DESeq2 (B). (A,B) Complete lists of disease-associated genes and significance are included in Supplementary tables. (A,B) 3 mice/group. (A) Horizontal bars indicate the mean, (B) mean ± S.E.M.

**Supplementary Figure 5 (Related to Figure 2).**
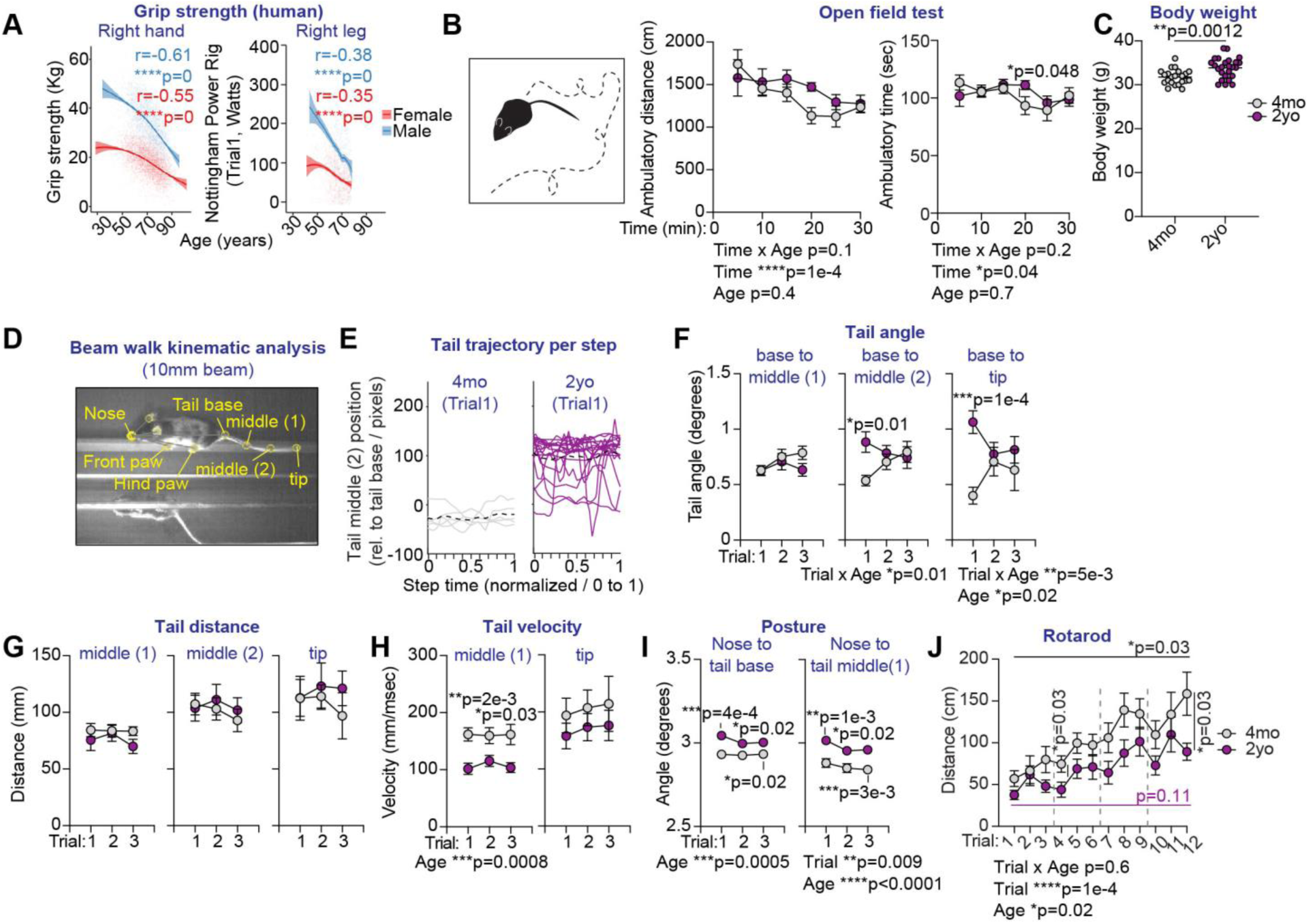
Additional characterization of motor deficits in aging. (A) Source data is publicly available through the Rancho San Bernardo Study of Healthy Aging and in a Supplementary Table. Correlation plots show individual data points and trend line indicating the relationship between grip strength of the right hand, and strength of the right leg with age. (B) Performance of 4mo (adult) and 2yo (aged) WT male mice in the open field test. (C) Body weight of 4mo (adult) and 2yo (aged) WT male mice. (D-I) Kinematic analysis of 4mo (adult) and 2yo (aged) WT male mice crossing the 10mm-wide beam. (D) Representative image shows the specific nodes that were video-tracked on the mouse body. (E) Representative tail trajectory traces indicate the position of the tail middle 2 segment relative to the tail base for every step taken by one 4mo (adult), and one 2yo (aged) mouse during trial 1 of this task. Black dashed line indicates average trajectory. (F) Tail angle of each tail segment relative to the tail base at the beginning of every step, (G) distance travelled by tail segments and (H) tail velocity at every step. (I) Angle between nose and tail segments at the beginning of every step indicate posture. (J) Distance travelled by 4mo (adult) and 2yo (aged) WT male mice on the accelerating rotarod. (A) Pearson correlation analysis, (B, F-J) RM 2-way ANOVA with Sidak’s multiple comparisons test, (C) Welch’s t-test. (A) Number of data points: 7,521 (hand), 1,691 (leg), (B) 16-19 mice/group, (C) 22-26 mice/group, (F-J) 10-13 mice/group. (C) Horizontal bars indicate the mean, (B,F-J) mean ± S.E.M.

**Supplementary Figure 6 (Related to Figure 3).**
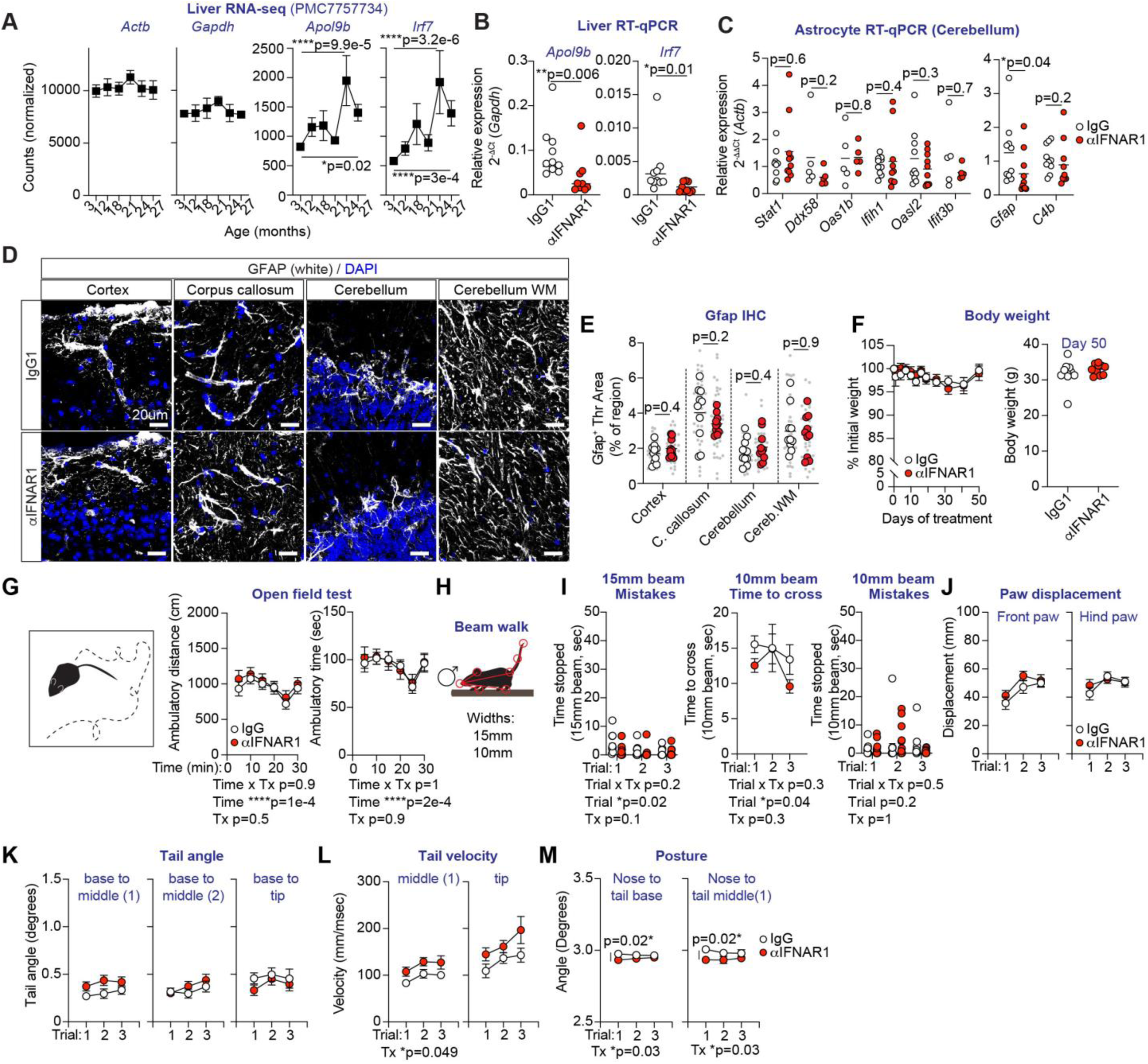
Additional characterization of type I IFN signaling promotes peripheral inflammation and motor deficits in aged mice. (A) Published RNA-seq data of the liver of WT male and female mice in adulthood and aging. Gene expression level in DESeq2 normalized counts at the indicated timepoints. *Actb* and *Gapdh* expression is stable through the lifespan, whereas *Apol9b* and *Irf7* increase with age. (B,C) 2yo (aged) WT male mice are intraperitoneally (i.p.) treated with IgG1 control or anti-IFNAR1 antibodies for 60 days. Expression level of *Apol9b* and *Irf7* normalized to *Gapdh*, in the liver of IgG1 and anti-IFNAR1 treated mice at the experimental endpoint. (C) RT-qPCR analysis of astrocyte nuclei in the cerebellum at the experimental endpoint using *Actb* as reference gene. (D) Gfap protein detected with immunohistochemistry in different brain regions in IgG1 and anti-IFNAR1 treated mice at the experimental endpoint. (E) Quantification of Gfap signal in cortex, corpus callosum, cerebellum cortex and cerebellum white matter. (F) Percentage of initial weight over time (left) and grams of body weight on day 50 of treatment (right). (G) Performance of mice in the open field test. (H-M) Mice are analyzed while crossing 15mm, and 10mm-wide elevated beams. (I) Time to cross the 10mm-wide beam, and time stopped on 15mm and 10mm-wide beams. (J) Paw displacement, (K) tail angle of each tail segment relative to the tail base at the beginning of every step, (L) tail velocity of indicated tail segments, and (M) angle between nose and tail segments at the beginning of every step. (A) False Discovery Rate (FDR) values obtained from web browser hosting data in PMC7757734. (B,C,E) Mann-Whitney tests, (F,G, I-M) RM 2-way ANOVA with Sidak’s multiple comparisons test. (A) 3-4 male mice/group, (B,E-M) 10 mice/group, (C) 5-10 mice/group. (B,C,E,F,I) Horizontal bars indicate the mean, (A,F,G,I,J-M) mean ± S.E.M.

**Supplementary Figure 7 (Related to Figure 3).**
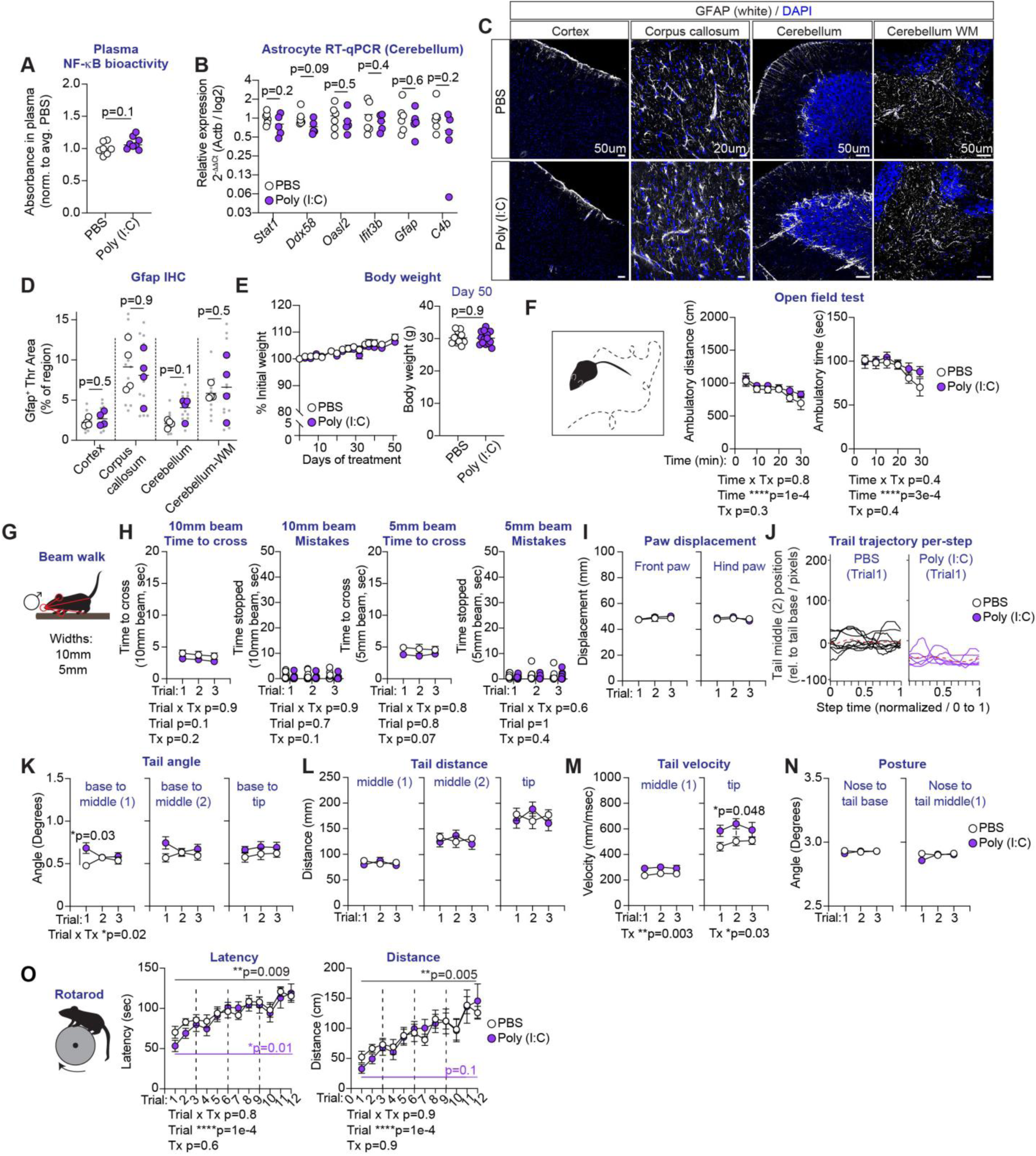
Peripheral low-level viral inflammation does not promote astrocyte antiviral state nor motor deficits. (A-O) 3mo (adult) WT male mice are i.p. treated with PBS or low dose Poly (I:C) (0.4 mg/kg) for 52 days. (A) NF-kB bioactivity in plasma 3hrs after the last injection. (B) RT-qPCR analysis of astrocyte nuclei in the cerebellum at the experimental endpoint using *Actb* as reference gene. (C) Gfap protein detected with immunohistochemistry in different brain regions in treated mice at the experimental endpoint. (D) Quantification of Gfap signal in cortex, corpus callosum, cerebellum cortex and cerebellum white matter. (E) Percentage of initial weight over time (left) and grams of body weight on day 50 after treatment (right). (F) Performance of mice in the open field test. (G-N) Mice are analyzed while crossing 10mm, and 5mm-wide beams. (H) Time to cross and time stopped while crossing 10mm and 5mm-wide beams. (I) Paw displacement. (J) Representative tail trajectory traces indicate the position of the tail middle 2 segment relative to the tail base for every step taken by one PBS, and one Poly (I:C)-treated mouse during trial 1 of this task. Red dotted line indicates average trajectory. (K) Angle of each tail segment relative to the tail base at the beginning of every step, (L) distance travelled by tail segments and (M) tail velocity for every step. (N) Angle between nose and tail segments at the beginning of every step indicates posture. (O) Latency and distance travelled on the accelerating rotarod. (A,B,D) Mann-Whitney tests, (E,F,H-O) RM 2-way ANOVA with Sidak’s multiple comparisons test. (A) 8 mice/group, (B) 5-6 mice/group, (C,D) 4 mice/group, (E-O) 13-14 mice/group. (A,B,D,E) Horizontal bar indicates the mean, (E,F,H,I,K-O) mean ± S.E.M.

**Supplementary Figure 8 (Related to Figure 4).**
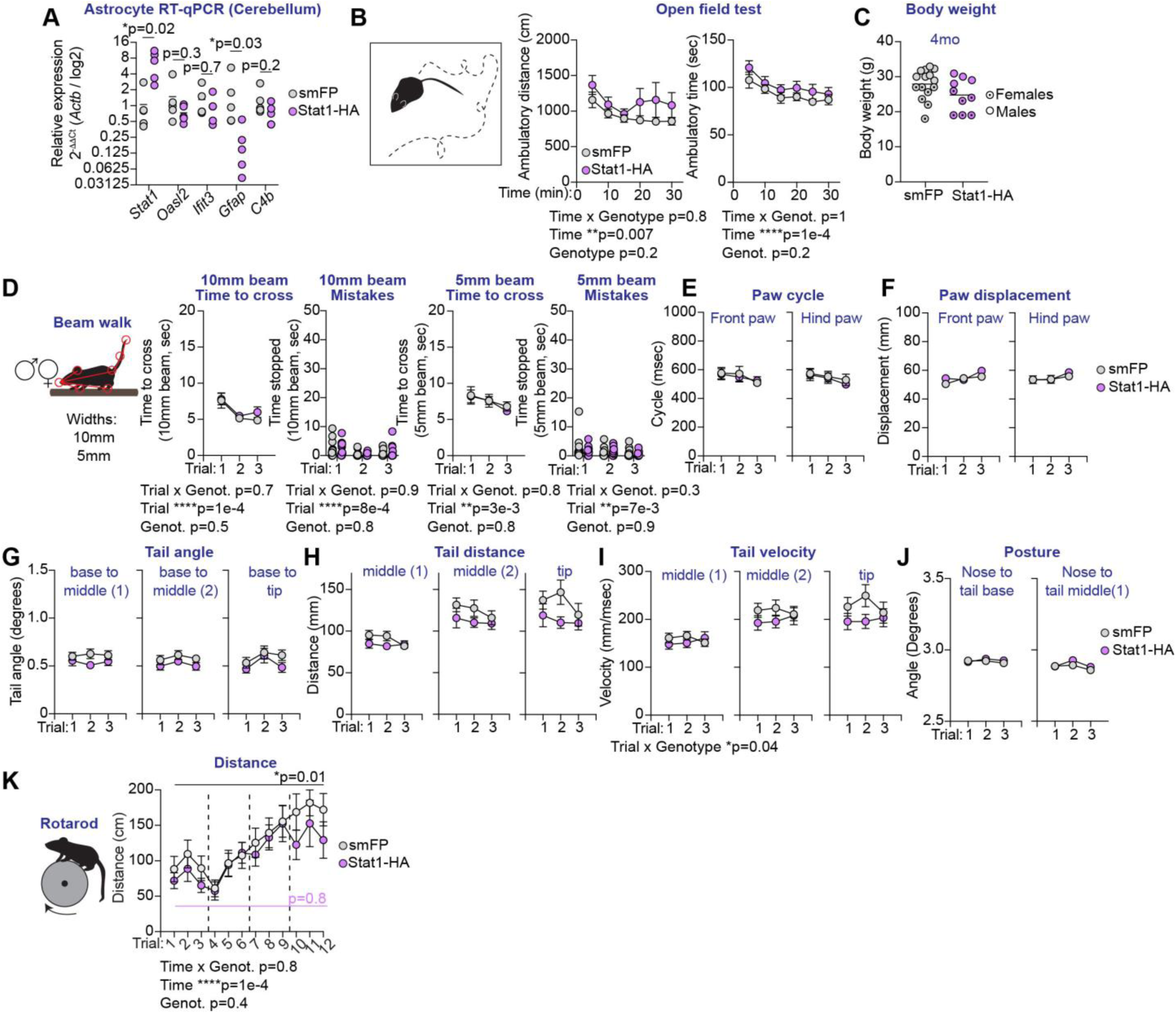
Astrocyte Stat1 does not promote astrocyte antiviral state or motor deficits in adult mice. (A-K) Male and female WT mice are r.o. injected AAVs at post-natal days 12-16. Behavioral testing is performed at 4mo, and the experiment terminated at 4.5mo. (A) RT-qPCR analysis of astrocyte nuclei in the cerebellum at the experimental endpoint using *Actb* as reference gene. (B) Performance of mice in the open field test. (C) Mouse body weight at 4mo. (D-J) Mice are analyzed while crossing 10mm, and 5mm-wide beams. (D) Time to cross and time stopped while crossing 10mm and 5mm-wide beams. (E) Paw cycle, (F) paw displacement, (G) angle of each tail segment relative to the tail base at the beginning of every step, (H) distance travelled by tail segments and (I) tail velocity for every step. (J) Angle between nose and tail segments at the beginning of every step indicates posture. (K) Distance travelled on the accelerating rotarod. (A) Multiple Mann-Whitney tests, (B, D-K) RM 2-way ANOVA with Sidak’s multiple comparisons test. (A) 5 mice/group, (B-K) 14-15 mice/group. (A,C,D) Horizontal bar indicates the mean, (B,D-K) mean ± S.E.M.

**Supplementary Figure 9 (Related to Figure 4).**
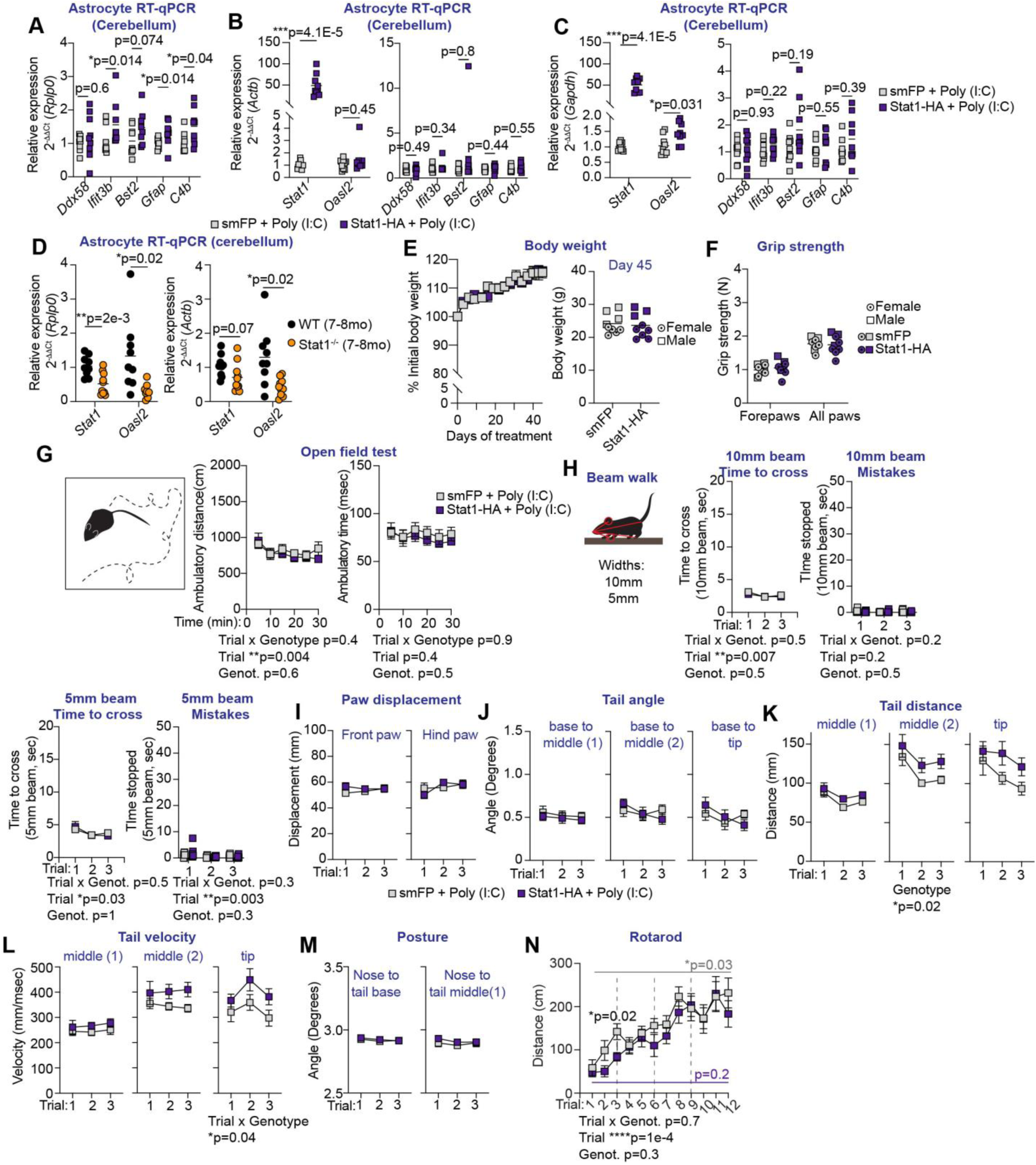
Additional characterization of astrocyte Stat1 promotes motor deficits during chronic inflammation. (A-C,E-N) Male and female WT mice are r.o. injected AAVs at post-natal days 12-16. Starting at 2mo, both groups of mice are i.p. treated with low dose Poly (I:C) (0.4 mg/kg) for 52 days. Behavioral testing is done between day 40 and 47 of treatment, and the experiment terminated at 4.5mo. (A-C) RT-qPCR analysis of astrocyte nuclei in the cerebellum at the experimental endpoint in smFP and Stat1-HA mice, or (D) at 7-8mo in WT and Stat1-/- male and female mice. Reference genes are *Rplp0* (A, D-left), *Actb* (B,D-right), and *Gapdh* (C). (E) Percentage of initial weight over time (left) and grams of body weight on day 45 of treatment (right). (F) Grip strength of fore limbs or all limbs. Circles and squares are the average of 3 independent trials per mouse. (G) Performance of mice in the open field test. (H-M) Mice are analyzed while crossing 10mm, and 5mm-wide beams. (H) Average time to cross, and time stopped on 10mm and 5mm-wide beams for every trial. (F) PCA using 45 kinematic parameters after analysis of the 5mm-wide beam. (I) Paw displacement, (J) angle of each tail segment relative to the tail base at the beginning of every step, (K) distance travelled by tail segments and (L) tail velocity for every step. (M) Angle between nose and tail segments at the beginning of every step indicates posture. (N) Distance travelled on the accelerating rotarod. (A-D) Multiple Mann-Whitney tests, (E,G-N) RM 2-way ANOVA or (F) 2-way ANOVA with Sidak’s multiple comparisons test. (A-N) 9 mice/group. (A-F) Horizontal bar indicates the mean, (E,G-N) mean ± S.E.M.

**FigureS10. Supplementary Figure 10 (Related to Figure 5).**
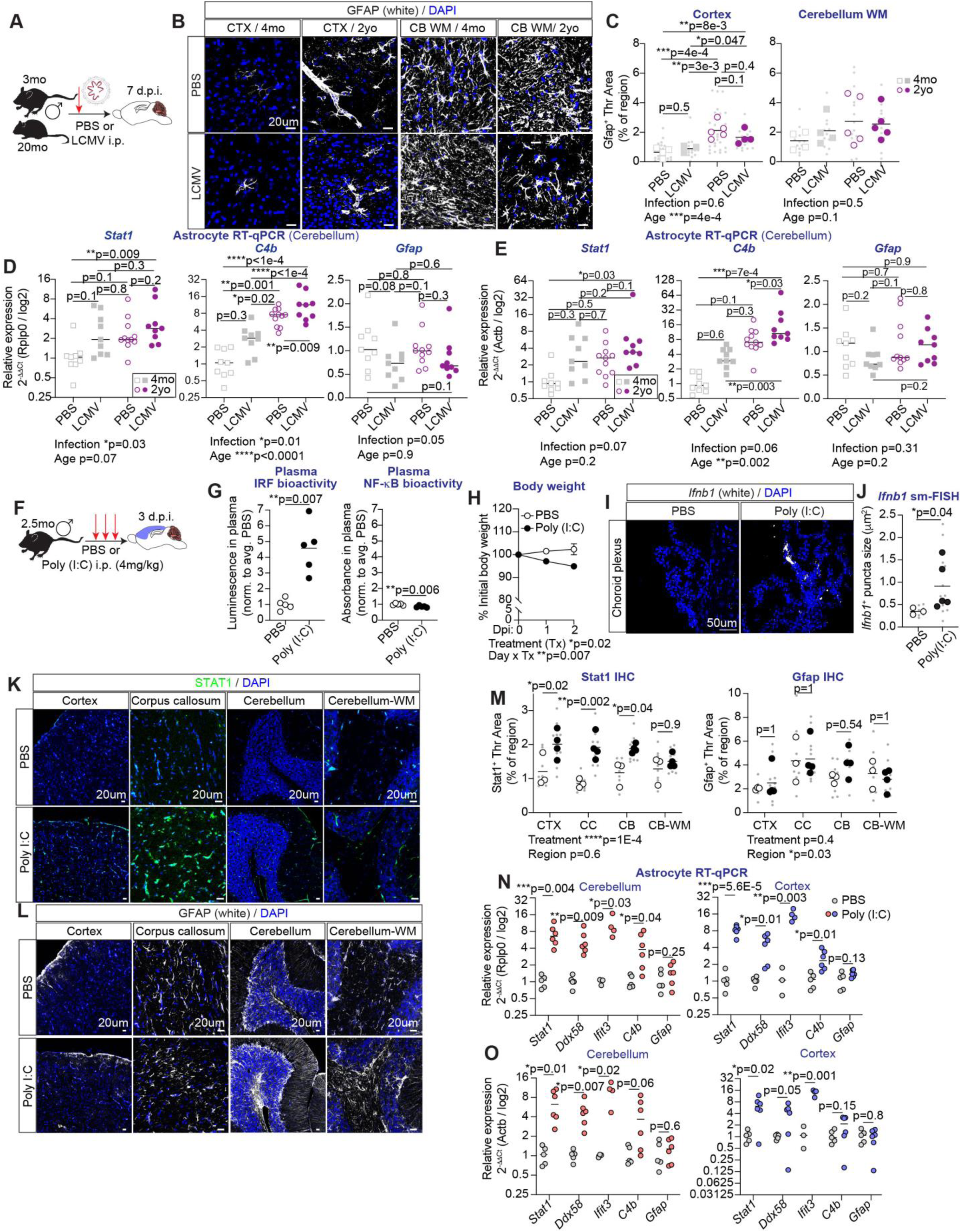
Systemic high-level viral inflammation promotes antiviral gene signatures in astrocytes in the cortex and cerebellum. (A-E) 3mo (adult) and 20mo. (aged) male mice are i.p. injected with PBS or 2x10^5^ plaque forming units (pfu) of Lymphocytic choriomeningitis virus Armstrong. Analyses are performed at day 7 post-injection (dpi). (B) Gfap protein detected with immunohistochemistry in the cortex (CTX) and cerebellar white matter (CB-WM) at indicated ages. (C) Quantification of Gfap signal in (B). Grey small dots indicate individual images, grey squares and magenta dots indicate individual mice. (D,E) RT-qPCR analysis of astrocyte nuclei in the cerebellum at the experimental endpoint using (D) *Rplp0* and (E) *Actb* as reference genes. (F-O) 2.5mo (adult) male mice are i.p. injected with PBS or high dose Poly (I:C) (4mg/kg) for 3 consecutive days. Analysis is done 3hrs after the last dose, on day 3 post-injection (dpi). (G) IRF and NF-kB bioactivity in plasma 3hrs after the last injection. (H) Percentage of initial weight over time. (I,J) *Ifnb1* mRNAs detected using sm-FISH in the choroid plexus in the lateral and 4^th^ ventricles of PBS and Poly (I:C) injected mice. (J) Quantification of *Ifnb1* puncta size in (I). (K-M) Stat1 (K) and Gfap (L) proteins detected with immunohistochemistry in cortex, corpus callosum, cerebellum cortex and cerebellum white matter (WM). (M) Quantification of Stat1 and Gfap signals in (K,L). Grey small dots indicate individual images, white and black circles indicate individual mice. (N,O) RT-qPCR analysis of astrocyte nuclei in the cortex and cerebellum at the experimental endpoint using (N) *Rplp0* and (O) *Actb* as reference genes. (C,D,E,M) 2-way ANOVA or (H) RM 2-way ANOVA with Sidak’s multiple comparisons test. (G,J,N,O) Mann-Whitney tests. (B,C) 4-5 mice/group, (D,E) 9-11 mice/group, (G) 5 mice/group, (H) 4 mice/group, (I,J,M) 3-5 mice/group, (N,O) 5-6 mice/group. (C,D,E,G,J,M,N,O) Horizontal bar indicates the mean, (H) mean ± S.E.M.

**Supplementary Figure 11 (Related to Figure 5).**
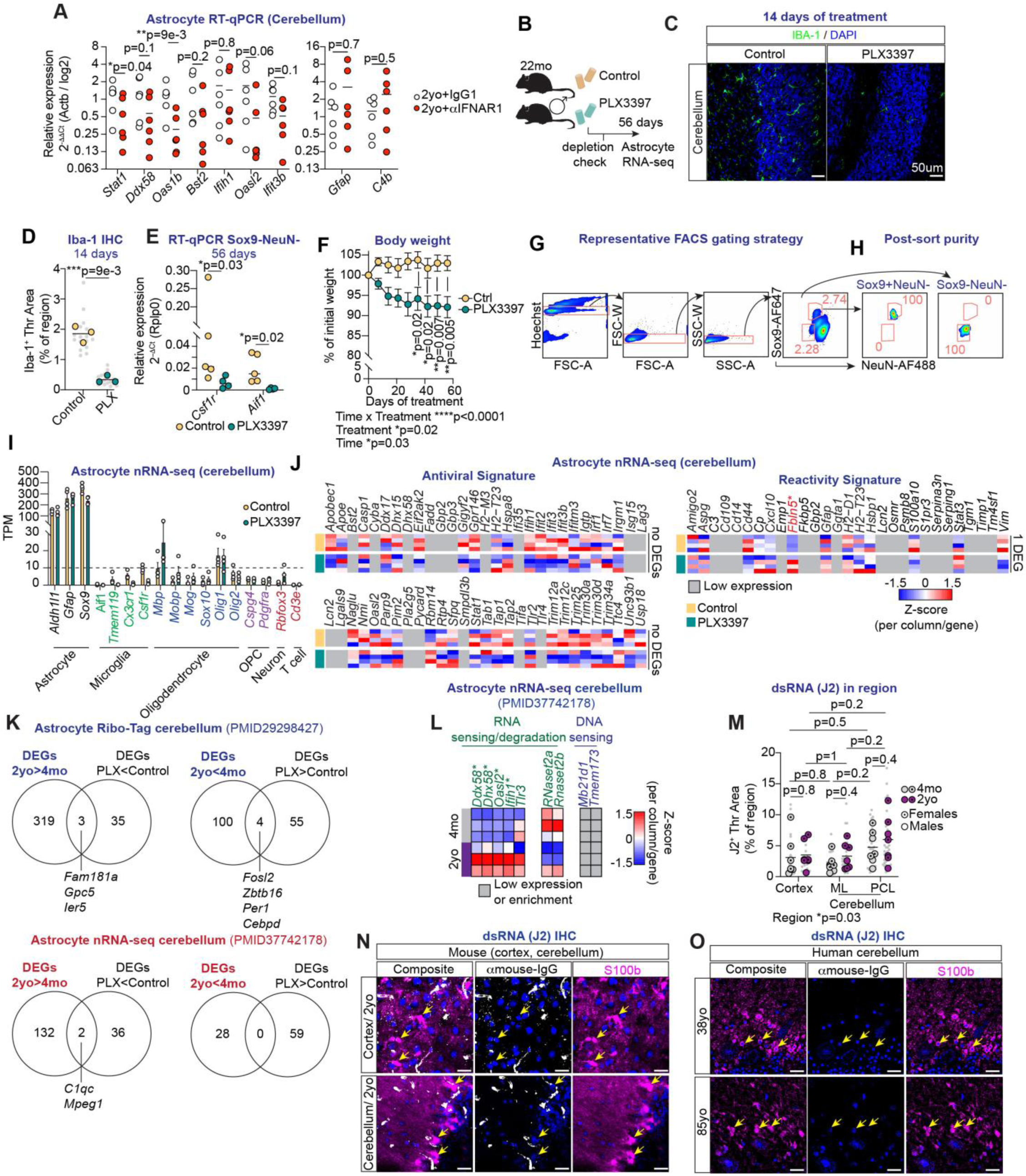
Additional characterization of local type I IFN signaling promotes astrocyte antiviral state in the aged cerebellum. (A) 20-22mo aged WT male mice are injected IgG1 or anti-IFNAR1 into the cerebellum. Cerebella are analyzed 48hrs after injection. RT-qPCR analysis of astrocyte nuclei in the cerebellum at the experimental endpoint using *Actb* as reference gene. (B-K) 22mo aged WT male mice are given control or PLX3397-formulated diet for 2 months. (C) Iba-1 protein is detected with immunohistochemistry in the cerebellum 14 days after initiation of treatment. (D) Quantification of Iba-1 signal in the cerebellum. Grey dots are individual images; yellow and green circles are mouse averages. (E) RT-qPCR of FACS-purified non-astrocyte, non-neuronal Sox9-NeuN- nuclei 56 days after initiation of treatment using *Rplp0* as reference gene. (F) Percentage of initial weight over time in control and PLX3397 mice. (G) Representative gating strategy to FACS-purify Sox9+NeuN- astrocyte nuclei, and Sox9-NeuN- non-astrocyte, non-neuron nuclei. Gating consists of placing rectangular gates around single nuclei using FSC-A vs. Hoechst, FSC-A vs FSC-W and SSC-A vs SSC-W. A polygon gate was drawn around Sox9+ NeuN-, or Sox9- NeuN- to gate on astrocyte or non-astrocyte, non-neuron nuclei, respectively. Gate numbers represent the percentage of single nuclei that are Sox9+ NeuN- or Sox9- NeuN-. (H) Representative post-sort purity for each population. (I) RNA-seq analysis of astrocyte nuclei in the cerebellum 56 days after initiation of treatment. An expression cut-off of 10 transcripts per million (TPM) is used, which enriches for astrocyte transcripts over microglia, oligodendrocyte, oligodendrocyte precursor cell (OPC), neuron, and T cell transcripts. (J) Heatmaps show Z-score normalized gene expression of antiviral and reactivity-associated genes in astrocytes in the cerebellum. Grey color indicates low expression. (K) Venn diagrams overlap PLX-dependent DEGs with DEGs in cerebellar astrocytes in 2yo (aged) vs 4mo (adult) mice obtained in Ribo-Tag and nRNA-seq datasets. (L) Heatmaps show Z-score normalized gene expression of genes involved in RNA sensing and degradation, and DNA sensing. Astrocyte nRNA-seq data in PMID37742178 was used. (M) Quantification of total double stranded (ds)RNAs using the J2 antibody, not restricted to S100b+ soma. Quantification of J2 signal in cortex, and cerebellar molecular layer (ML) and Purkinje Cell layer (PCL). Grey small dots indicate individual images, grey and magenta circles indicate individual mice. (N,O) Staining controls using secondary antibody in the absence of J2 primary antibodies in mouse (N) and human samples (O). (A) 6 mice/group, (B-K) 4 mice/group, (M) 7 mice/group. (A,D,M) Horizontal bar indicates the mean, (F,I) mean ± S.E.M.

**Supplementary Figure 12 (Related to Figure 5).**
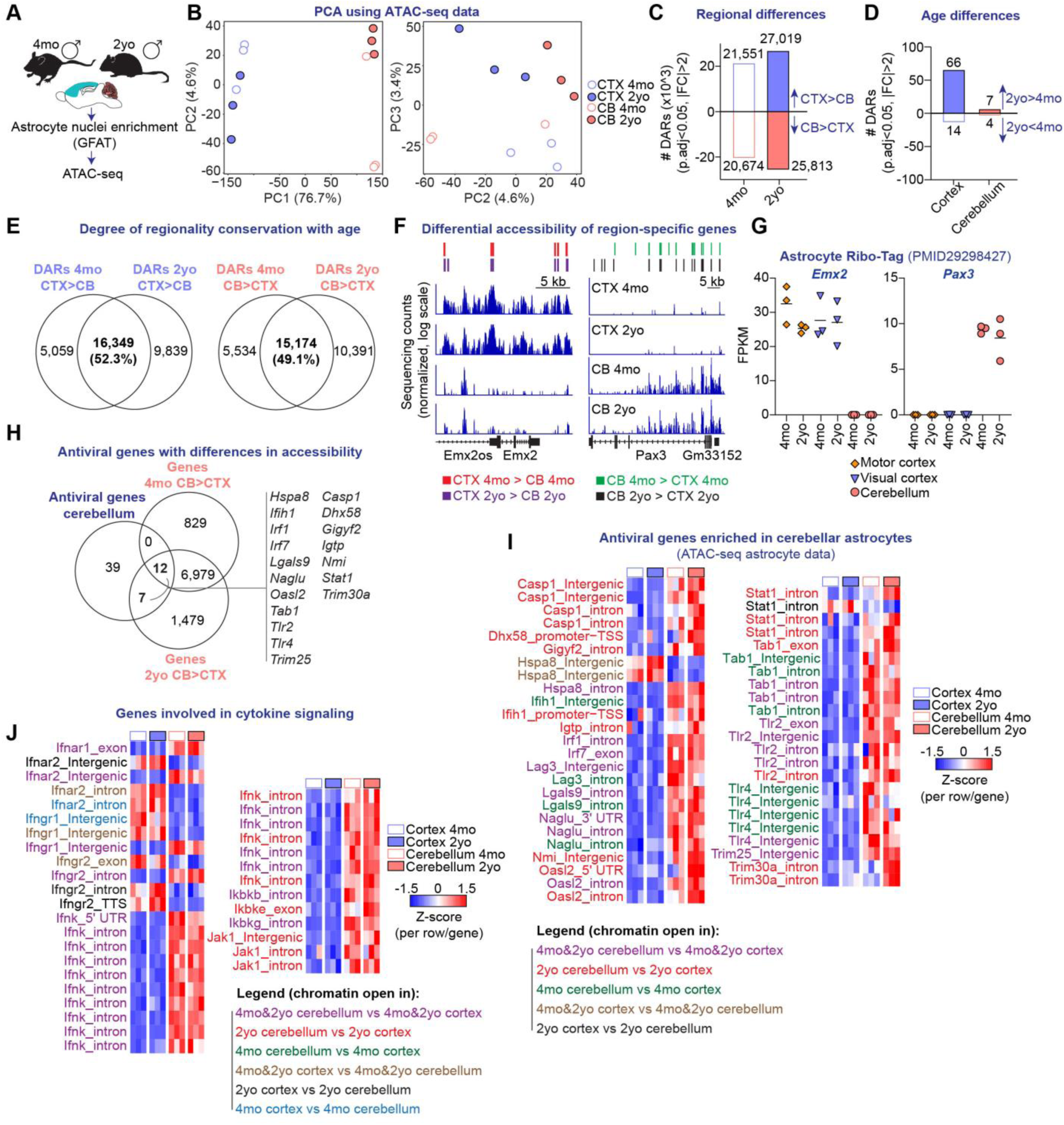
Regional differences in the chromatin landscape of cortical and cerebellar astrocytes in adulthood and in aging. (A-F, H-J) Astrocyte nuclei in the cortex and cerebellum of 4mo and 2yo male WT mice (3 mice/group) are enriched using Glyoxal Fixed Astrocyte Transcriptomics followed by Assay for Transposase-Accessible Chromatin using sequencing (ATAC-seq). (B) PCA using all chromatin accessible regions. Axes indicate PC coordinates, and percentages are variability explained by each component. Astrocyte nuclei in cortex (CTX), and cerebellum (CB) at different ages are shown. Circles represent individual mice. (C) Differentially accessible chromatin regions (DARs) between cortex and cerebellum in adulthood and in aging. This indicates regional differences in chromatin accessibility at 4mo and 2yo. (D) Age differences in chromatin accessibility in cortical and cerebellar astrocytes. DARs are defined with p.adj<0.05, and absolute fold change (FC) over 2. (C,D) DARs include accessible regions in the promoter and transcriptional start site (TSS), 3’ untranslated regions, exons, intergenic regions, introns, non-coding regions, and transcriptional termination site. (E) Venn diagrams overlap regionally accessible regions at 4mo and 2yo. (F) Representative DNA library tracks with differentially accessible regulatory regions for *Emx2* and *Pax3* by region. Specific comparisons are color coded. Representative tracks for each gene use the same scale, allowing visual comparison across regions and ages. (G) Expression level of *Emx2* and *Pax3* in region-specific astrocytes in 4mo and 2yo mice determined using Ribo-Tag. (H) Venn diagram identifies 19 antiviral genes with differential accessible regions in astrocytes in the cerebellum, compared to cortex, at 4mo and 2yo. (I,J) Heatmaps show Z-scores using normalized tag counts for each individual sample at indicated sites, for antiviral genes that were differentially accessible, and identified in H (I), and for differentially accessible genes involved in cytokine signaling (J). (I,J) The specific comparisons at which differences in accessibility are color-coded, and are specified in the legend. (G) Horizontal bar indicates the mean.

## Methods

### Mice

All mouse work was approved by The Salk Institute Institutional Animal Care and Use Committee (IACUC). Mice were housed in the Salk Institute Animal Resources Department on a light cycle of 12 hours of light and 12 hours of dark, with *ad libitum* access to food and water.

#### WT mice

Experiments use C57BL/6 male and female mice. Adult (2.5-4mo) mice were obtained from the Jackson Laboratories or from in-house breeders. Mice at post-natal days 12-16 were obtained from in-house breeder pairs. Aged (20-24mo) mice were obtained from the National Institutes of Aging (NIA).

#### Transgenic mouse lines

Ifnar1-/- and Stat1-/- retired breeder mice were a generous gift from Dr John Teijaro at The Scripps Research Institute. Mice were originally obtained from the Jackson Laboratories (RRID:IMSR_JAX:028288 & RRID:IMSR_JAX:012606).

### Mouse treatments

Whenever possible, mice treated with control or experimental compounds were co-housed to minimize potential cage effects in behavior or astrocyte phenotype. Experiments using control or PLX3397-formulated diets are an exception.

#### *In vivo* treatments with Poly (I:C)

Polyinosinic-polycytidylic acid sodium salt (Poly (I:C), Tocris#4287) was diluted in sterile PBS1x at a concentration of 10 mg/mL, aliquoted and stored at -80C. On the day of injection, mice were weighed, and aliquots were thawed and diluted in sterile PBS1x at a specified concentration in a final volume of 100 uL per mouse.

##### Chronic treatment with low-dose Poly (I:C)

Adult (2.5-3mo) mice were intraperitoneally (i.p.) injected with 0.4 mg/kg Poly (I:C) in 100uL or sterile PBS1x every 3^rd^ day for up to 60 days. Behavior analysis was started on day 40 after initiation of treatment. Behavioral assays were performed on days when mice were not injected or, if on an injection day, they were injected after behavior concluded. At endpoint, mice were collected 3 hrs after the last dose was administered.

##### Acute treatment with high-dose Poly (I:C)

Adult (2.5-3mo) mice were i.p. injected with 4 mg/kg Poly (I:C) in 100uL or sterile PBS1x every 24hrs for 3 consecutive days. Mice were collected 3 hrs after the last dose was administered.

#### *In vivo* antibody treatments

##### Peripheral IFNAR-1 blockade

Aged (20-22mo) mice were i.p. treated with IgG1 (MOPC-21, BioXCell) or IFNAR-1 (MAR1-5A3, BioXCell) antibodies. Antibody stocks (10 mg/mL) were kept sterile at 4C, and were diluted in sterile PBS1x the day of the injection. An initial dose of 500 ug i.p. was administered on day 0, followed by 250 ug i.p. every 3^rd^ day, for a total of 60 days. Behavioral experiments were performed starting on day 40 post-treatment and through endpoint. Behavioral assays were performed on days when mice were not injected or they were injected after behavior concluded. Mice were collected 18 hours after the last dose was administered.

##### Cerebellar IFNAR-1 blockade

Aged (20-22mo) mice underwent stereotaxic surgery as explained in the ‘Method details’ section, and were injected 20ug in 2uL of IgG1 (MOPC-21, BioXCell) or IFNAR-1 (MAR1-5A3, BioXCell) antibodies into the cerebellum. Mice were collected 48 hrs after injection.

#### PLX3397 treatment

Rodent diet AIN-76A (Research Diets #D10001i) was formulated with 290mg PLX3397 (MedChem Express #HY-16749) per kg by Research Diets. Control AIN-76A and formulated diets were irradiated, and provided ad libitum. 22mo WT male mice were given control or PLX3397-formulated diets for a total of 56 days.

### Viral injections

#### AAV inoculation

Mice were anesthetized using 1.5-2% isoflurane, and retro-orbitally (r.o.) injected at post-natal day 12 to 16 (P12-16) with 1x10^11^ genome copies of AAVs of the PhP.eB serotype^1^.

#### Viral infection

Mice were intraperitoneally injected with 2x10^5^ plaque forming units of Lymphocytic choriomeningitis virus Armstrong^2^. Viral stocks were grown and quantified with standard plaque assay^3^, and were generously provided by Dr Susan Kaech. Uninfected and infected mice were housed separately.

### Cell cultures

#### Interferon regulatory factor (IRF) and NF-kB bioactivity assay

Raw-Dual Cells (InvivoGen, #rawd-ismip) report activity of interferon regulatory factors (IRFs) by secreting luciferase and NF-kB activity by secreting embryonic alkaline phosphatase (SEAP).

##### Bioactivity in brain lysates

*1) Tissue preparation:* Mice were anesthetized and underwent whole-body perfusion with PBS, followed by dissection of the cerebellum and the cortex, which was detached from sub-cortical areas and visible white matter was removed. Brain regions were snap frozen in liquid nitrogen and kept at -80C. Frozen tissues were thawed on ice, weighed, bead homogenized in 200ul of DMEM media (ThermoFisher #11965118) and spun down. Supernatant was aliquoted and frozen at -80C until the day of the assay. *2) Cell line preparation & assay*: Cells were cultured as indicated in the manufacturer’s protocol, and used for this assay after 4 to 8 passages. Each sample consisted of 20ul of brain homogenate, which was co-cultured with Raw-DUAL macrophages in a tissue culture incubator at 37C, 5% CO2 for 18 hours in testing media, as specified in the manufacturer’s protocol. 2-3 wells containing media were used as negative control. Positive control wells included media supplemented with 100 ng/ml of LPS (positive control for IRF and SEAP, Sigma #L2630) or 100 ug/ml Poly (I:C) (positive control for IRF, Tocris#4287). 18 hours after, 20uL of culture supernatant was collected for each sample, incubated with SEAP detection reagent and signal detected 6 hrs after. Luciferase signal in supernatant was read 24 hours after initiation of co-culture. Absorbance and luminescence signals were collected using an Infinite 200 Pro Microplate Reader (Tecan). *3) Data analysis*: Average luminescence or absorbance in negative control wells was subracted from raw signal in every biological sample. In each assay, we also confirmed signals over background in positive control wells. We divided each corrected luminescence or absorbance value by each sample’s weight, leading to luminescence or absorbance per mg of tissue.

##### Bioactivity in plasma

*1) Plasma collection:* Blood was drawn from the mouse submandibular facial vein using Goldenrod 5mm lancets (Braintree, NC9416572), and collected using heparin-coated glass capillaries (Caraway Micro Blood Collecting Tubes Red, Fisherbrand #0266825). Blood was transferred into 1.5 mL Eppendorf tubes, and centrifuged at 10,000 rpm for 10 min at 4C. Plasma was aliquoted and stored at -80C. *2) Cell line preparation & assay:* Cells were cultured as indicated in the manufacturer’s protocol, and used for this assay after 4 to 8 passages. Each sample consisted of 20ul of thawed plasma, which was co-cultured with Raw-DUAL macrophages in a tissue culture incubator at 37C, 5% CO2 for 18 hours in testing media, as specified in the manufacturer’s protocol. Given that plasma proteins can interfere with luciferase assays^4^, wells containing plasma from naïve, uninjected mice were used as negative control. Positive control wells included plasma from mice 3 hrs after injection with 12 mg/kg Poly (I:C), and in vitro controls 100 ng/ml of LPS, or 100 ug/ml Poly (I:C) in media. 20uL of culture supernatant was collected for each sample, incubated with SEAP detection reagent and signal detected 6 hrs after. Luciferase signal in supernatant was read 24 hours after initiation of co-culture. *3) Data analysis*: Average luminescence or absorbance in negative control wells was extracted from raw signal in every biological sample. Corrected values were normalized to the average of the control group (PBS) for each experimental cohort. We also checked increased signal of positive controls over negative control.

### Method details

#### Injection of antibodies into the cerebellum of aged mice using stereotaxic surgery

Mice were administered pre-operative subcutaneous (s.c.) carprofen (5 mg/kg), dexamethasone i.p. (2 mg/kg) and were anesthetized using 1.5-2% isoflurane. 6 sites were drilled on the skull above the cerebellum, at stereotaxic coordinates relative to bregma: −6.5 anterior posterior (AP), +/- 0.5 medial-lateral (ML); −7.5 AP, +/-1.8 ML; −8.0 AP, +/-1.4 ML. Antibodies were injected at these 6 locations at 3 depths relative to the surface of the brain (Z −2.5, −2.0, and −1.5). 2uL of antibody were distributed across the 18 injection sites (i.e. 111nl per site), and injected at a speed of 10nL/sec. The pipette was kept at each location for 3 minutes after each injection to allow diffusion, before moving to the next location. After injections, mice were sutured, administered saline s.c. (20 mL/kg) and placed back in their home cage on top of a heated blanket until awoken. Mice were s.c. administered carprofen (5 mg/kg) 24 hrs after surgery, and were collected 48 hrs after surgery.

#### Brain collection

Mice were anesthetized with intraperitoneal injection of a mixture of 100mg/kg ketamine (Victor Medical Company 1699053) and 20mg/kg xylazine (Victor Medical Company 1264078) until unresponsive to toe pinch.

##### Fresh tissue collection

was performed for experiments involving RNA-seq or ATAC-seq assays, and consisted of brain dissection immediately after anesthesia and cervical dislocation.

##### Fixed tissue collection for IHC

Mice were anesthetized and underwent whole-body perfusion with room temperature PBS and 4% PFA paraformaldehyde solution (PFA) (Electron Microscopy Sciences 19210), followed by brain collection. Brains were placed in 4% PFA solution at 4C for 18-20 hrs. After that, brains were washed 3 times for 5min in PBS1x, were submerged in 30% sucrose for 72 hrs, after which they were embedded in Tissue Freezing Medium (Electron Microscopy Sciences 72593), frozen in a dry ice/ethanol slurry and stored at −80C.

##### Fixed tissue collection for experiments involving astrocyte RT-qPCR and IHC

Brains were collected immediately after anesthesia and cervical dislocation, without perfusion. Fresh brains were cut through the midline and one hemisphere was submerged in 4% PFA at 4C for 18-20 hrs, while the remaining hemisphere was used for astrocyte nuclei isolation and RT-qPCR analysis. After overnight fixation, brain hemispheres were washed 3 times for 5min in PBS1x, submerged in 30% sucrose for 72 hrs, embedded and frozen in Tissue Freezing Medium, and stored at −80C.

#### Viral vector generation

Glycerol stocks were expanded in an over-night culture in Luria Broth (LB) supplemented with carbenicillin (100 ug/mL) at 37C with shaking at 150rpm. DNA was purified using the endotoxin-free Qiagen Plasmid Maxi kit (Qiagen #12162) following manufacturer’s instructions. Plasmids were sequenced using Plasmidsaurus before packaging into adeno-associated virus particles (AAV) at the Salk Institute Viral Vector (GT3) core. Plasmids were packaged into the AAV PHP.eB serotype with titers of 3.84E14 genome copies (gc)/mL (GfaABC1D-smFP, control protein) and 4.36E14 gc/mL (GfaABC1D-Stat1-HA, Stat1 plasmid). The GfaABC1D-Stat1-HA plasmid was generated in-house, and GfaABC1D-smFP was used as in^5^.

#### Generation of GfaABC1D-Stat1-HA plasmid

We obtained a modified coding sequence for mouse Stat1-HA cDNA in a backbone containing an origin of replication and ampicillin resistance gene from Twist biosciences (Ensembl CCDS ID: CCDS78581, Transcript ID: ENSMUST00000186057). To increase the efficiency of protein translation, a Kozak sequence was inserted before the start codon, and the stop codon was replaced with the sequence of a hemagglutinin (HA) tag. The modified Stat1 CDS was inserted into a pZac2.1 GfaABC1d-tdTomato vector (Addgene #44332) using the in-Fusion HD cloning kit (Takara 638909) following manufacturer’s instructions. First, the GfaABC1d-tdTomato vector was linearized using NotI-HF and NheI-HF restriction enzymes (New England Biolabs, R3189S, NheI-HF) for 18 hrs at 37C. To avoid plasmid re-ligation, 5’ ends were de-phosphorylated using recombinant Shrimp Alkaline Phosphatase (rSAP NEB #M0371, 1U/ug of DNA) for 30min at 37C followed by enzyme inactivation at 65C for 5min. Linearized and de-phosphorylated plasmid was purified from an agarose gel and cleaned up using Qiaquick PCR & Gel Cleanup kit (Qiagen #28506) following manufacturer’s instructions. Stat1 CDS was PCR-amplified from its original plasmid using Q5 High-Fidelity 2x Master Mix (NEB M0492S) and primers designed with in-Fusion Cloning Primer design tool (Takara) (Forward 5’ to 3’: CTCACTATAGGCTAGCGCCACCATGTCACAGTGGT; Reverse 5’ to 3’: CTGCTCGAAGCGGCCGCCTAAGCGTAATCTGGAACATCGT). Size of PCR product was evaluated on an agarose gel, and cleaned up on a column using Qiaquick PCR & Gel Cleanup kit (Qiagen #28506). The In-Fusion HD cloning enzyme premix was used to insert Stat1 CDS into the previously linearized vector, leading to the GfaABC1D-Stat1-HA plasmid. 3uL of the in-Fusion reaction were used to transform 50uL of One-Shot Stbl3 Chemically Competent E. coli (ThermoFisher C737303) according to the manufacturer’s protocol. Colonies were screened on agarose plates containing carbenicillin (100 ug/mL; Teknova C2130). Individual colonies were expanded in an over-night culture in Luria Broth (LB) supplemented with carbenicillin (100 ug/mL), and aliquoted 1:500 into 250 mL of LB carbenicillin, which was cultured for 18-20 hrs at 37C with shaking at 150rpm. DNA was purified using the endotoxin-free Qiagen Plasmid Maxi kit (Qiagen #12162) following manufacturer’s instructions. Plasmids were sequenced using Plasmidsaurus before packaging into AAVs.

#### RT-qPCR analysis of the mouse liver

2.5x2.5cm of liver (left lobe) were harvested, placed in Eppendorf tubes at −80C. 1) *Liver RNA extraction:* tubes containing tissues were brought to −20C inside a cryostat and were sectioned into pieces of 30mg. 30mg of frozen liver per mouse were placed in tubes containing 600uL of RLT solution (Qiagen) supplemented with beta-mercaptoethanol (10uL/mL of RLT) and with 0.5mL of 1mm zirconia disruption beads (Research Products International). Tissues were homogenized using a BeadRuptor 12 homogenizer (Omni International), centrifuged at 5,000 rpm for 5 min at 4C, and supernatant collected. Supernatants were passed through Qiashredder and gDNA eliminator columns before RNA extraction using the RNeasy Mini Kit (Qiagen #74106) following manufacturer’s instructions. RNA was eluted in 30uL of molecular grade water pre-warmed at 65C. 2) *RT-qPCR*: RNA concentration was estimated using nanodrop. 1ug of RNA per sample were used in a 20uL RT reaction using SuperScript IV Vilo MasterMix (Invitrogen #11756050). QPCR was ran in 5uL reactions using 0.4uL cDNA, 5ul Sybr Green PCR Master Mix (Applied Biosystems #4309155), 0.28uL forward primer (10uM), 0.28uL reverse primer (10uM), and 4.04ul of molecular grade water. We ran duplicate reactions using a QuantStudio Real-Time PCR system (ThermoFisher). Cycling conditions were as follows: 1) 95C for 10min, 2) 95C for 15sec (denaturation), 3) 60C for 1min (annealing and extension), 4) steps 2 and 3 for 40 cycles, 5) 95C for 15sec (melt curve stage 1), 6) 60C for 1min using ramp −1.6C/sec (melt curve stage 2), 7) 95C for 15sec using ramp 0.075C/sec (dissociation stage). For each biological sample, cycle threshold (Ct) values were normalized to Actb or Gapdh following the 2-dCt method. Primers sequences used (all 5’ to 3’): Actb-fw CATTGCTGACAGGATGCAGAAGG, Actb-rev TGCTGGAAGGTGGACAGTGAGG, Gapdh-fw CATCACTGCCACCCAGAAGACTG, Gapdh-rev ATGCCAGTGAGCTTCCCGTTCAG, Apol9b-fw GGAAGTCTGGTGCTCTCAGCAA, Apol9b-rev ATGAGTGCCAGTCAGAGCAGCT, Irf7-fw CCTCTGCTTTCTAGTGATGCCG, Irf7-rev CGTAAACACGGTCTTGCTCCTG.

### Immunohistochemistry (IHC) procedures

#### Mouse brain cryosectioning

Sagittal brain sections of 18-20 μm were obtained from all mouse cohorts using a cryostat (Hacker Industries OTF5000). 3 tissue sections per slide were cut from fixed frozen brains at −20C, and placed on Superfrost Plus micro slides (VWR 48311-703). Slides were dried for 1 hr at room temperature and stored at −80C for future use. Slides were used within 6 months after cutting.

#### Stat1 detection in the aged brain

18-20um-thick fixed sagittal brain sections were cryosectioned from fixed brains at −20C, and stored at −80C until use. Fixed-perfused brains were used for this particular stain. 1X Epitope retrieval solution was made by diluting 10X RNAscope Target Retrieval solution (ACD #322001) in molecular grade water (Thermo Scientific #A57775), and was pre-warmed at 90C for 30 min. Dried slides were retrieved from −80C, immersed in PBS1x for 2 min, placed in 1X epitope retrieval solution, and incubated at 90C for 20 min. After that, slides were rinsed in PBS1x for 1 min, placed in 100% ethanol (Fisher scientific #BP2818100) for 2 min, and dried at room temperature for 10 min. A hydrophobic barrier was drawn around sections and slides were dried for 10 additional minutes. Slides were blocked and permeabilized in blocking buffer (10% goat serum in PBS with 0.3% Triton) for 1 hr at room temperature. Primary antibodies were diluted in antibody buffer (5% goat serum in PBS with 0.3% Triton), and stained for 48 hrs in a humified chamber at 4C. We used rabbit Stat1 (D1K9Y clone, CST #14994, 1:100) and guinea pig S100b (Polyclonal, Synaptic systems #287004, 1:500). After staining, slices were washed 3 times in PBS1x for 10 min. Secondary antibodies were diluted in antibody buffer and slices incubated for 2 hrs at room temperature. Secondary antibodies used were goat anti-rabbit AlexaFluor488 (Invitrogen A-11008,1:500), goat anti-guinea pig AlexaFluor647 (Invitrogen A-21450, 1:500). After staining, slides were washed 3 times for 10 min in PBS1x, followed by DAPI staining (ThermoFisher #D1306, 1:10,000) for 5 min. To quench lipofuscin autofluorescence, we used TrueBlack Lipofuscin Autofluorescence Quencher 20X in DMF (Biotum #23007). TrueBlack was diluted in 70% ethanol (15uL TrueBlack in 300 uL 70% ethanol), added to slides and incubated for 5min at room temperature. Slides were washed 3 times for 10 min in PBS1x, and coverslips were mounted using Fluoromount-G Mounting Medium (ThermoFisher #00-4958-02).

#### Additional stains in the mouse brain

18-20um-thick fixed sagittal brain sections were washed in PBS1x for 2min, air dried at room temperature, a hydrophobic barrier drawn around them, and additional drying. Slides were blocked and permeabilized in blocking buffer (10% goat serum in PBS with 0.3% Triton) for 1 hr at room temperature. Primary antibodies were diluted in antibody buffer (5% goat serum in PBS with 0.3% Triton), and stained for 48 hrs in a humified chamber at 4C. Primary antibodies used with this protocol are mouse J2 (D1K9Y clone, CST #14994, 1:100), rat Gfap (Polyclonal, Synaptic systems #287004, 1:500), rabbit Iba-1 (FujiFilm Biosciences #016-20001, 1:500), rabbit S100b (EP1576Y clone, Abcam #52642, 1:500), guinea pig S100b (Polyclonal, Synaptic Systems #287004, 1:500), rat HA (3F10 clone, Sigma #11867423001, 1:250). To detect Stat1 after Poly (I:C) injection, rabbit Stat1 D1K9Y clone, CST #14994, 1:100) was used. After staining, slices were washed 3 times in PBS1x for 10 min. Secondary antibodies were diluted in antibody buffer and slices incubated for 2 hrs at room temperature. Secondary antibodies used were goat anti-rabbit AlexaFluor488 (Invitrogen A-11008,1:500), goat anti-guinea pig AlexaFluor647 (Invitrogen A-21450, 1:500). After staining, slides were washed 3 times for 10 min in PBS1x, followed by DAPI staining (ThermoFisher #D1306, 1:10,000) for 5 min. To quench lipofuscin autofluorescence, we used TrueBlack Lipofuscin Autofluorescence Quencher 20X in DMF (Biotum #23007). TrueBlack was diluted in 70% ethanol (15uL TrueBlack in 300 uL 70% ethanol), added to slides and incubated for 5min at room temperature. Slides were washed 3 times for 10 min in PBS1x, and coverslips were mounted using Fluoromount-G Mounting Medium (ThermoFisher #00-4958-02).

#### Human cerebellum staining

1) *Paraffin embedding:* Post-mortem, formalin-fixed human cerebellar vermis from de-identified adult and elderly human individuals were obtained from the NIH NeuroBioBank. Specimens originated from male Caucasian individuals with no brain pathology including dementias or other neurological diseases. Adult ages were 34, 38 and 41 years old, and elderly were 80, 81, 85, and 88 years old. Tissues were received in formalin, embedded in paraffin and sectioned into 10um-thick sections in the Tissue Technology Shared Resource at the Moore’s Cancer Center (UC San Diego). 2) *Deparaffination*: tissue sections were baked at 60C for 35 min followed by 2 de-waxing washes of 5 min each in 100% xylene (Sigma #214736). Sections were rehydrated by placing them in decreasing amounts of ethanol (100, 95, 50, 20 % ethanol in molecular grade water, 2 min for each incubation). After the last ethanol step, sections were washed in PBS1x for 2min, and in PBS 0.1% tween for 5 min and 3 times. *3) Permeabilization*: PBS1x + 0.3% Triton was added to sections for 10 min at room temperature. *4) Background removal*: To quench lipofuscin autofluorescence, we used TrueBlack Lipofuscin Autofluorescence Quencher 20X in DMF (Biotum #23007). TrueBlack was diluted in 70% ethanol (15uL TrueBlack in 300 uL 70% ethanol), added to slides and incubated for 5min at room temperature. Slides were washed 3 times for 10 min in PBS1x. Subsequent steps are performed in the absence of detergents. *5) Antigen retrieval:* 1X Epitope retrieval solution was made by diluting 10X RNAscope Target Retrieval solution (ACD #322001) in molecular grade water (Thermo Scientific #A57775), and was pre-warmed at 90C for 30 min. Slides were placed in 1X epitope retrieval solution, and incubated at 90C for 10 min. After that, slides were rinsed in PBS1x for 1 min and a hydrophobic barrier was drawn around them. *6) Blocking*: Blocking buffer (5% goat serum, 2% BSA in PBS1x) was added to slides for 1 hr at room temperature. *7) Antibody staining*: Primary antibodies were diluted in blocking buffer and incubated for 48 hrs in a humified chamber at 4C. Antibodies used are guinea pig S100b Polyclonal, Synaptic systems #287004, 1:500) and mouse J2 (D1K9Y clone, CST #14994, 1:100). After staining, slices were washed 3 times in PBS1x for 10 min. Secondary antibodies were diluted in blocking buffer and slices incubated for 2 hrs at room temperature. Secondary antibodies used were goat anti-mouse AlexaFluor555 (Invitrogen #A21422,1:500), goat anti-guinea pig AlexaFluor647 (Invitrogen A-21450, 1:500). After staining, slides were washed 3 times for 10 min in PBS1x, followed by DAPI staining (ThermoFisher #D1306, 1:10,000) for 5 min. Coverslips were mounted using Fluoromount-G Mounting Medium (ThermoFisher #00-4958-02).

#### Single molecule Fluorescent in situ hybridization (sm-FISH) in mouse brain

RNAscope V2 (ACD Bio-techne) reagents were used. 18-20um-thick fixed sagittal brain sections were retrieved from −80C freezer, placed into PBS1x for 5 min at room temperature to remove TFM, and baked at 60C for 30 min. Slides were post-fixed in 4% PFA (Electron Microscopy Sciences 19210) at 4C for 15 min. 1) *Tissue dehydration*: slides were placed in ethanol/water solutions with increasing ethanol content for 5 min at room temperature for each concentration (50% ethanol in molecular grade water, 70%, and 100%). Slides were dried in 100% ethanol (Fisher scientific #BP2818100) at −20C for 18-20 hrs. Slides were removed from ethanol, dried for 10 min at room temperature, hydrophobic barrier drawn around them, and added RNAscope Hydrogen Peroxide for 10 min at room temperature. Slides were washed in molecular grade water 3 times for 2 min. 2) *Protease treatment*: slides were incubated in Protease III for 10 min at 40C. 3) *Probe hybridization*: hybridization and amplification steps were performed as described in the manufacturer’s protocol. The following probes were used: Mm-Stat1-C1 (#479611), negative control probe-DapB-C2 (#310043), Mm-Ifnb1-C1 (#406531). Opal 570 dye was used for C1 probes (1:1000), and Opal 690 for C2 (1:1000). 4) *Counterstain, background quenching & mounting*: after the HRP step in the RNAscope assay, slides were blocked and permeabilized in PBS1x + 0.3% Triton for 1hr at room temperature. Primary antibodies were diluted in antibody buffer (5% goat serum in PBS with 0.3% Triton), and stained for 48 hrs in a humified chamber at 4C. Primary antibodies used with this protocol are rabbit S100b EP1576Y clone, Abcam #52642, 1:500) and guinea pig S100b (Polyclonal, Synaptic Systems #287004, 1:500). After staining, slices were washed 3 times in PBS1x for 10 min. Secondary antibodies were diluted in antibody buffer and slices incubated for 2 hrs at room temperature. Secondary antibodies used were goat anti-rabbit AlexaFluor488 (Invitrogen A-11008,1:500) or goat anti-guinea pig AlexaFluor647 (Invitrogen A-21450, 1:500). After staining, slides were washed 3 times for 10 min in PBS1x, followed by DAPI staining (ThermoFisher #D1306, 1:10,000) for 5 min. To quench lipofuscin autofluorescence, we used TrueBlack Lipofuscin Autofluorescence Quencher 20X in DMF (Biotum #23007). TrueBlack was diluted in 70% ethanol (15uL TrueBlack in 300 uL 70% ethanol), added to slides and incubated for 5min at room temperature. Slides were washed 3 times for 10 min in PBS1x, and coverslips were mounted using Fluoromount-G Mounting Medium (ThermoFisher #00-4958-02).

#### Image acquisition

For visualization of protein stains and RNAscope assays, tissue sections were imaged using LSM700 or LSM710 confocal microscopes (Zeiss) at 20X magnification as 8-bit images of 1-4 tiles, and as a Z-stack that imaged a total of 4.5um of thickness, in 4 slices of 1.50 um centered in the center of each section. Scanning speed was set to 6-7 (i.e. scan time ranging from 15.49 sec to 30.98 sec), with bi-directional scanning, by frame, and averaging of 2 images (using the mean) was used. In experiments requiring co-localization analysis, we routinely chose channels with the smallest overlapping spectra possible. For each region imaged and stain we measured 3-6 sections per mouse or human specimen.

#### Image analysis

Signal quantification was performed using macros in ImageJ. Each individual image was converted to 8-bit, a maximum intensity projection image using all stacks was created, and channels were split. The channel containing signal of interest was chosen, and an automatic Threshold was set using the ‘Triangle’ or ‘Otsu’ methods. The most suitable method was chosen after careful inspection of thresholded images, the criteria was to use the one that captured the signal of interest and discriminated background the most. Images were converted to mask, then saved in .tif format. To quantify signal within regions of interest (ROI): ROIs were manually drawn either delimiting whole regions (e.g. cerebellar layers), or around individual astrocyte soma based on S100b+ staining. ROIs were applied to their corresponding thresholded images, and % of thresholded area occupied by signal within each ROI was recorded. For quantification of *Ifnb1* signal in the choroid plexus, ROIs marking the lateral and 4^th^ ventricle choroid plexus were manually drawn. ROIs were overlaid to their corresponding thresholded image containing *Ifnb1* signal, and mean particle size within each ROI was obtained using ‘Analyze Particles’. Image analysis of human J2 staining was performed using automatic thresholding with the ‘Triangle’ method with the signal range clipped to 16-255 on the J2 channel. This range was established to eliminate background signal, and was determined based on negative staining control samples stained using only the secondary antibody in the J2 channel.

### Astrocyte transcriptomics and chromatin accessibility

#### Sample collection

Mice were anesthetized with intraperitoneal injection of a mixture of 100mg/kg ketamine (Victor Medical Company 1699053) and 20mg/kg xylazine (Victor Medical Company 1264078) until unresponsive to toe pinch. <u>For snRNA-seq:</u> Cortices and cerebella from adult (4mo) or aged (20-22mo) male mice (2 mice/group) were collected and pooled, yielding a total of 4 samples (cortex 4mo, cerebellum 4mo, cortex 2yo, cerebellum 2yo). <u>For ATAC-seq:</u> Cortices and cerebella from adult (4mo) or aged (20-22mo) male mice (3 mice/group) were collected and processed independently, yielding a total of 12 samples (3 cortex 4mo, 3 cerebellum 4mo, 3 cortex 2yo, 3 cerebellum 2yo).

#### Astrocyte nuclei isolation

Astrocyte nuclei were obtained using Glyoxal Fixed Astrocyte Transcriptomics (GFAT)^6^. Samples were placed in ice-cold PBS1x right after collection. When all samples were collected, cortices and cerebella were chopped using a blade and into fragments of approximately 9 mm^3^ in size. Fragments were submerged in 4mL of ice-cold glyoxal acidic solution (3% w/v glyoxal [Millipore Sigma 128465], 20% ethanol [Fisher BP2818100], 0.75% acetic acid [Millipore Sigma A6283], 5mM NaOH [Millipore Sigma 72068], in ddH2O) for 20 min at 4C. Samples were washed 3 times in PBS1x and placed in a dounce homogenizer containing ice-cold 3 mL NIMT buffer (in mM: 250 sucrose, 25 KCl, 5 MgCl2, 10 Tris-Cl pH 8, 1 DTT; 1:100 dilution of 10% Triton) supplemented with protease inhibitor (Roche mini tabs #11836153001, 1 tab for 11mL NIMT), 20U/mL of SUPERaseIN (Invitrogen #AM2694), 40 U/mL RNaseOUT (Invitrogen #10777019). Tissue was manually homogenized using the two-step Dounce homogenizer system (A and B) (Sigma #D9063) and placed on ice. Homogenates were mixed with 2 mL 50% iodixanol (OptiPrep Density Gradient Medium; Sigma #D1556) and underlaid with 2.5 mL of 25% iodixanol to create a gradient. Samples were centrifuged in a swinging bucket rotor (Sorval HS-4) at 10,000 xg for 20 min at 4C. Nuclei pellets were re-suspended in 1 mL ice-cold DPBS supplemented with 1% w/v BSA (DPBS+BSA), 20U/mL SUPERaseIN, 40U/mL RNaseOUT. Nuclei were stained for 5 min using 5 uM Hoechst 33342 solution, followed by centrifugation at 1,200 rpm for 5min at 4C. Pellets were stained in 100uL DPBS+BSA containing mouse anti-NeuN-AlexaFluor488 (Millipore #MAB377X; 1:1000) and rabbit anti-Sox9 antibody (Abcam AB185966; 1:100) for 1hr on ice. Nuclei were washed by adding 300 uL of DPBS+BSA, followed by centrifugation at 1,200 rpm for 5min at 4C. Pellets were stained in 100uL DPBS+BSA containing secondary antibody goat anti-rabbit IgG-AlexaFluor647 (Invitrogen #A-21224; 1:200) for 1hr on ice. Nuclei were washed in 300 uL DPBS+BSA, centrifuged at 1,200 rpm for 5min at 4C and re-suspended in 500ul DPBS+BSA.

#### Fluorescence activated cell sorting

FACS experiments used a BD FACS Aria Fusion sorter located in the Salk Institute Flow Cytometry core. 70-100um nozzles were used to sort astrocyte nuclei. Hoechst-positive nuclei were gated, followed by doublet exclusion using forward and side scatter parameters (FSC-A vs SSC-A, and FSC-W vs SSC-W). To select the astrocyte population, a gate was placed in nuclei with positive Sox9 and devoid of NeuN signals (Sox9+NeuN-). Astrocyte nuclei were sorted into 1.5mL Eppendorf tubes pre-coated and filled with 300uL of DPBS+BSA. The number of Sox9+NeuN- astrocyte nuclei collected per sample depends on the downstream assay; 50,000 for single-nucleus (sn)RNA-seq, 150,000-200,000 nuclei for bulk nucleus (n)RNA-seq, 25,000-50,000 nuclei for RT-qPCR, and 30,000 nuclei for ATAC-seq. To check post-sort purity, we pooled 5uL of each sorted sample, ran it through the FACS and checked that >95% of nuclei fell within the Sox9+NeuN- astrocyte gate. After sorting, tubes containing nuclei were centrifuged at 1,200 rpm for 5min at 4C, and pellets used in downstream assays (snRNAseq, bulk nRNAseq or RT-qPCR). FACS acquisition was performed using BD FACSDiva software, and representative gating strategy was done using FlowJo software v10.10.0.

#### RT-qPCR of astrocyte nuclei

After sorting, nuclei pellets were resuspended in 350 uL of RLT buffer (Qiagen RNeasy Micro kit#74004) supplemented with beta-mercaptoethanol (10 uL/ml; Sigma M3148) and stored at - 80C. 1) *RNA extraction:* RNA was extracted after thawing lysates at room temperature, with the RNeasy micro kit (Qiagen #74004) following manufacturer’s instructions with on-column DNase digestion, and elution in 14 uL of water pre-warmed at 65C. 2) *RT-qPCR*: RNA abundance was below the limit of detection for nanodrop or Qubit assays, as we have previously reported ^6^. 7uL per sample were used in a 20uL RT reaction using SuperScript IV Vilo MasterMix (Invitrogen #11756050). QPCR was ran in 5uL reactions using 0.4uL cDNA, 5ul Sybr Green PCR Master Mix (Applied Biosystems #4309155), 0.28uL forward primer (10uM), 0.28uL reverse primer (10uM), and 4.04ul of molecular grade water. We ran duplicate reactions using a QuantStudio Real-Time PCR system (ThermoFisher). Cycling conditions were as follows: 1) 95C for 10min, 2) 95C for 15sec (denaturation), 3) 60C for 1min (annealing and extension), 4) steps 2 and 3 for 40 cycles, 5) 95C for 15sec (melt curve stage 1), 6) 60C for 1min using ramp −1.6C/sec (melt curve stage 2), 7) 95C for 15sec using ramp 0.075C/sec (dissociation stage). For each biological sample, cycle threshold (Ct) values were double normalized to Rplp0, Actb or Gapdh and to the average Ct of the control group following the 2-ddCt method. Primers sequences used (all 5’ to 3’): Rplp0-fw GCTTCGTGTTCACCAAGGAGGA, Rplp0-rev GTCCTAGACCAGTGTTCTGAGC; Actb-fw CATTGCTGACAGGATGCAGAAGG, Actb-rev TGCTGGAAGGTGGACAGTGAGG; Gapdh-fw CATCACTGCCACCCAGAAGACTG, Gapdh-rev ATGCCAGTGAGCTTCCCGTTCAG; Stat1-fw GCCTCTCATTGTCACCGAAGAAC, Stat1-rev TGGCTGACGTTGGAGATCACCA; Ddx58-fw AGCCAAGGATGTCTCCGAGGAA, Ddx58-rev ACACTGAGCACGCTTTGTGGAC; Oas1b-fw CTGTGCTGACCTCAGAGAAGTC, Oas1b-rev TGCCCTTGAGTGTGGTGCCTTT; Ifih1-fw TGCGGAAGTTGGAGTCAAAGCG, Ifih1-rev CACCGTCGTAGCGATAAGCAGA; Oasl2-fw CCAAAACGAGGTCGTCAGGAAC, Oasl2-rev AGCCACCTGTTCCCATCCCTTT; Ifit3b-fw GCTCAGGCTTACGTTGACAAGG, Ifit3b-rev CTTTAGGCGTGTCCATCCTTCC; Gfap-fw CACCTACAGGAAATTGCTGGAGG, Gfap-rev CCACGATGTTCCTCTTGAGGTG; C4b-fw GGAGAGTGGAACCTGTAGACAG, C4b-rev CACTCGAACACGAGTTGGCTTG; Ifit3-fw GCTCAGGCTTACGTTGACAAGG, Ifit3-rev CTTTAGGCGTGTCCATCCTTCC; Bst2-fw CAAACTCCTGCAACCTGACCGT, Bst2-rev CTCCTGGTTCAGCTTCGTGACT; Csf1r-fw TGGATGCCTGTGAATGGCTCTG, Csf1r-rev GTGGGTGTCATTCCAAACCTGC; Aif1-fw TCTGCCGTCCAAACTTGAAGCC, Aif1-rev CTCTTCAGCTCTAGGTGGGTCT. Primers used only in Stat1-/- mice: Stat1(exon5)-fw ACGCTGCCTATGATGTCTCG, Stat1(exon5)-rev AGAAAAGCGGCTGTACTGGT.

#### Bulk nRNA-sequencing of astrocyte nuclei after PLX3397 treatment

After sorting, nuclei pellets were resuspended in 350 uL of RLT buffer and RNA was extracted exactly as explained in the RT-qPCR of astrocyte nuclei section. RNA quality and concentration were estimated using TapeStation High-sensitivity RNA (Agilent). Libraries were constructed using 9ng of RNA per sample with the Illumina Stranded Total RNA prep (with Ribo-Zero Plus) sequencing kit, following manufacturer’s instructions. Libraries were sequenced at 2.5–3.5 × 10^7^ reads per sample, paired end 100 bp reads using a NovaSeq S4 sequencer. Quality control of sequenced reads was done using FastQC, mapping to the Mus musculus genome (mm10) with STAR v2.5.3a^7^. To count number of reads per gene, we used ‘MakeTagDirectory’ and ‘analyzeRepeats’ in HOMER^8^ with the ‘-count genes’ option to include reads mapped to introns and exons. Counts were exported in raw format, and in transcripts per million (TPM). Differential expression analysis was performed using DESeq2 v1.46.0^9^. Differentially expressed genes (DEGs) were selected using the following criteria: Benjamini-Hochberg (BH) adjusted p-value < 0.05, absolute fold change (FC) > 1.5, and average expression cut-off of equal or above 10 TPM for the group with highest expression in the comparison. An expression cut-off of 10 TPM was chosen to avoid transcripts expressed by non-astrocytes.

#### snRNA-seq of astrocyte nuclei

*1) Single-nuclei assay:* Tubes containing 50,000 FACS-purified astrocyte nuclei per sample were centrifuged at 1,200 rpm for 5min at 4C, re-suspended in 43.3 uL of DPBS+BSA, and loaded onto a Next GEM Chip G included in the 10X Chromium Next GEM Single Cell 3’ v3.1 kit (10X Genomics). Barcoding, cDNA generation and library preparation were performed using the manufacturer’s protocol with no modifications. cDNA and library concentration, and quality were estimated using Qubit Fluorimeter (Thermo Fisher) and Tape Station (Agilent), respectively. Libraries were sequenced using NovaSeq S1 at The Salk Institute Next Generation Sequencing Core at ∼50,000 reads/cell. Sequencing of each sample was followed by alignment to the mouse genome (mm10), filtering, barcode counting, and Unique Molecular Identifier (UMI) counting with Cell Ranger v8.0.0^10^ (‘cellranger count’ function with the ‘include-introns’) followed by aggregation of all 4 datasets (i.e. 4mo cortex and cerebellum, and 2yo cortex and cerebellum) with the ‘cellranger aggr’ function. *2) Data analysis*: all downstream analyses were performed in R with Seurat 5.3.0^11^. Cells with 500 to 5,000 detected genes (nFeature_RNA > 500 and < 5,000) and with less than 5% of mitochondrial RNA (percent.mt < 5) were kept for downstream analyses. Normalization and variance stabilization was done with the ‘SCT transform’ function^12^ (SCT v2), followed by non-linear dimensionality reduction with Uniform Manifold Approximation and Projection (UMAP)^13^. We used MapMyCells (Allen Institute for Brain Science) using Correlation clustering, which detected a small fraction of neurons, microglia and oligodendrocytes that were excluded from downstream analyses. The average number of UMIs per astrocyte (nCount_RNA) was 4,697 ± 2,663, and average number of genes (nFeature_RNA) was 1,853 ± 629. We next performed dimensionality reduction and cluster identification using ‘RunPCA’, ‘RunUMAP’ (with dimensions 1:40), ‘FindNeighbors’ (with dimensions 1:40), and ‘FindClusters’ (with resolution 0.1). Metadata columns were added to the Seurat object to identify samples by age, and brain region. Data were sub-set into two independent Seurat objects (one for adult and aged cortical astrocytes, and one for cerebellar). ‘RunPCA’, ‘RunUMAP’, ‘FindNeighbors’ and ‘FindClusters’ were re-run for each object using aforementioned parameters (and cluster resolution 0.1). To determine cluster markers, we used the ‘FindAllMarkers’ function (wilcox test), and selected only genes that were positively enriched in the cluster being analyzed compared to all others, with adjusted p.value<0.05, and expression in at least 15% of cells (pct>0.15). Differentially-expressed genes between 4mo and 2yo samples, for each cluster, were determined using the ‘FindMarkers’ function (wilcox test). Enrichment for antiviral, and reactivity-associated genes were calculated using ‘AddModuleScore_UCell’ with gene lists containing antiviral and reactivity-associated genes in Figure 1. To determine layer-enriched astrocyte sub-sets in the cortex, ‘AddModuleScore_UCell’ was used to calculate gene scores for genes enriched in astrocytes in upper, middle, deep and white matter cortical layers^14^. Specific genes included in each signature are included as a supplementary table. Histograms were created depicting the distribution of UCell scores for each layer gene set, for all astrocytes in the cortical dataset. DimPlots of all cortical astrocytes were created whereby the 25% of astrocytes with highest UCell scores for each layer gene signature were colored in red.

#### ATAC-seq of astrocyte nuclei

ATAC-seq was performed as previously described^15^. Following sorting, 30,000 nuclei were washed with cold PBS and were pelleted by centrifugation at 500g for 10 min at 4°C then resuspended in 50 ul transposition mix (25ul 2x TD buffer, 2.5 ul transposase (100 nM final), 16.5 ul PBS, 0.5 ul 1% digitonin, 0.5 ul 10% Tween-20, 5 ul H2O) and incubated at 37°C for 30 min in a thermomixer with 1000 RPM mixing. DNA was purified using a Qiagen MinElute PCR cleanup kit, then PCR amplified using indexed oligos. The optimal number of amplification cycles for each sample was determined by qPCR. Libraries were purified and size-selected with AMPure XP beads and sequenced on an Illumina NovaSeq X platform to generate 50 bp paired-end reads. Paired-end reads were aligned to the M. musculus mm10 genome using STAR^7^ with default parameters. ATAC-seq peaks were called using the HOMER (v4.11)^8^ findPeaks program using parameters for DNAse-seq (-style dnase). Differentially accessible regions were identified using DESeq2^9^ via the HOMER script *getDifferentialPeaksReplicates.pl*, applying thresholds of fold change ≥ 2.0 or ≤ −2.0 and FDR < 0.05. Peaks were annotated using *annotatePeaks.pl*, and signal tracks were generated from *bedGraphToBigWig* with default parameters for visualization. Unique peak IDs were assigned based on genomic coordinates using the *GenomicRanges* package (v1.60.0)^16^ within R (v4.5.0), ensuring consistent feature matching between read count tables and differential peak analyses.

#### Pathway analysis using published astrocyte transcriptomic datasets

Published astrocyte RNA-seq or MERFISH datasets were used^6,17–19^. *1) Differentially expressed gene (DEG) selection:* To identify DEGs in Labarta-Bajo, et al^6^ we used DESeq2 v1.46.0 to compare cerebellar astrocytes in 4mo vs 2yo WT male mice. DEGs were selected using the following criteria: Benjamini-Hochberg (BH) adjusted p-value < 0.05, absolute fold change (FC) > 1.5 and an expression cut-off of at least 5 TPM for the group average. To identify DEGs in Boisvert, et al^17^, we used significantly up-regulated DEGs provided in the original study and used an additional cut-off of absolute FC > 1.5. DEGs in Clarke et al^18^ were used as provided in the original study. *2) Pathway analysis:* DEGs that were up-regulated in aged compared to adult astrocytes were selected from each study, as explained above. We used enrichGO with ontology ‘Biological process’ included in the ‘clusterProfiler’ library in R. Pathways were enriched if at least 10 DEGs were present in the gene ontology term. *3) Interferon-related gene enrichment:* enrichment analysis for Interferon-related genes in astrocytes in different brain regions throughout the lifespan was performed using spatial transcriptomics MERFISH data in Sun et al^19^. A Seurat object was created from the provided .h5ad matrix, and astrocytes were sub-set using metadata annotations provided in the original study. We created a gene set that included Interferon-related genes in the MERFISH assay (*Ifitm1, Ifitm3, Bst2, Ifi27, B2m, Jak1, Ifit1, Ifnar2, H2-D1, H2-K1, and Stat1*). ‘AddModuleScore_UCell’ was used to compute enrichment scores for each astrocyte in the dataset. Average UCell scores for astrocytes in each brain region and time-point were calculated and plotted.

#### Astrocyte antiviral gene signature

To create an antiviral gene signature relevant to astrocytes in the aged cerebellum, we first merged all genes included in GO terms ‘Regulation of immune response’, ‘Response to virus’, and ‘Defense response to virus’, which were the top 3 enriched pathways in when using up-regulated DEGs in aged vs adult cerebellar astrocytes obtained with Ribo-Tag (Figure 1A&^17^). We further filtered out genes that were not DEGs, which yielded a list of 58 genes (Supplementary Table).

### Mouse behavior assays

Mice undergoing behavioral testing were acclimated to the behavior testing room for 1 hour prior to testing. Behavioral experiments were performed during the light cycle, between 12pm and 6pm. All mice were handled in the days preceding the beginning of testing, to allow habituation to experimenter.

#### Open field testing

Testing room was lit by 400 lux of indirect lighting through acclimation and testing. Open field boxes with dimensions of 43.2 cm x 43.2 cm x 30.5 cm height were used (ENV-515S-A, Med Associates). Boxes contain locomotor beams 0.5 cm off the ground, and rearing beams 6.5 cm beams off the ground. Individual mice were placed at the center of the arena, and allowed to explore for 30 min. Ambulatory distance and time exploring the whole arena were obtained in 5min intervals using Activity Monitor Software (v SOF-812, Med Associates).

#### Rotarod

Testing room was lit by 200 lux of indirect lighting through acclimation and testing. Testing was done using a Rotor-Rod apparatus 33’’ height x 36’’ width x 24’’ depth contained inside a black opaque box (SD Instruments). 4 individual mouse lanes were 4.25’’ wide, with fall height of 18’’, rod diameter of 1.25’’ and 7 photobeams per lane (28 total) spaced by 0.5’’. *1) Habituation of aged mice:* experiments comparing 4mo (adult) and 2yo (aged) mice involved a 3-day habituation period preceding the first testing day. For every habituation day, mice were placed on the static elevated rod for 30 seconds, followed by rotation of the rod at constant low speed (3RPM) for 60 sec. 3 trials were performed, each separated by at least 10min. Testing started the day after that. *2) Habituation of adult mice:* 2.5-4mo mice were habituated during the first day of testing. Mice were placed on the static elevated rod for 30 seconds followed by 60 sec at 3RPM for 3 trials. Testing proceeded right after that. *3) Testing of all mice:* for every trial, mice were placed on the static rod for 30 sec, followed by accelerated rod rotation from 0-30RPM in 300sec. 3 trials were performed, each separated by at least 10min. Time before falling off the rod (latency) and distance ran on the rod (distance) were recorded using Rotor-Rod software. 4 testing days were performed, spaced every 24-48hrs.

#### Grip strength

Testing room was lit by 200 lux of indirect lighting through acclimation and testing. A grip strength apparatus (Conduct Science) was used. Mice were gently picked up from the tail and allowed to hang on to the wire plate with fore paws, or with fore and hind paws. Mouse body was maintained horizontal, at a flat angle before gently pulling from the tail base until they let go. Peak force was recorded in newton (N) for 3 trials for each limb group. Average peak force among 3 trials is reported.

#### Beam walk

Testing room was lit by 30 lux of indirect lighting through acclimation and testing. A lamp was attached to recording camera to improve contrast. A beam walk set-up was built in-house. It consists of 2 elevated platforms (0.5m above the ground), each ending in a small enclosed shelter (‘home box’) to encourage mouse crossing. Beams of varying widths (30, 15, 10, and 5mm) and fixed length (1m) connect the two platforms. A mirror positioned at a 45^-^degree angle directly below the beam allows visualization of paw placement, and a side camera (Basler ace #acA2040) recorded mice undertaking the task at a rate of 30 frames per second. Mice crossed beams of reducing width in subsequent days, with 30, 15 and 10mm for experiments involving 2yo aged mice, or 30, 15, 10, and 5 mm for experiments using 2.5-4mo adult mice. Prior to recording the first trial, mice were placed in the middle of the beam to stimulate crossing toward either ‘home box’ at the edges. Mice were then recorded on their left side while crossing from the right toward the left edge. Mice were re-placed in the right home box, and allowed to cross 3 times. Mice completed 3 consecutive trials per session, and testing was repeated every 24 hrs. Video visualization using ImageJ/Fiji was used to determine the time required to cross a 67-cm segment of the beam, marked with stickers for reference. Mice that lost balance or made significant errors often paused on the beam. Thus, the total time spent stopped was recorded and used as a proxy for mistake rate.

#### Kinematic analysis of beam walk

Social LEAP Estimates Animal Poses (SLEAP) v 1.4.1^20^ was used to track parts of the mouse body while crossing narrow beams. Nose, ears, fore and hind left paws, tail base and 3 tail segments were tracked by manually labeling 1,000-3,000 frames per experiment, and training using a multi-animal bottom-up model. Trained models were applied to all videos, and their outputs were visually inspected for accuracy. When errors were detected, additional frames were annotated and models re-trained until predictions were optimal. 2-dimensional coordinates for every limb across time were exported in .h5 format. We used MATLAB (MathWorks) to calculate locomotor parameters for every trial, and data were exported in .csv format. For every trial, the first 2 and last 2 steps were eliminated, and the average for all steps within a trial were calculated. Kinematic data in this manuscript are mouse averages for every trial. Values for all kinematic parameters and statistical testing used in this study are available in Supplementary tables. SLEAP models, videos, and MATLAB code are available in <u>10.5281/zenodo.17477698</u>.

#### Human behavior data analysis

De-identified demographic and physical ability data are publicly accessible through The Rancho San Bernardo Study of Healthy aging^21^. Graphing and statistics were performed with R software v4.4.2 and Rstudio v2023.6.2.561. ‘Ggplot’ was used to plot grip and leg strength or time to cross a narrow line, against age, in male and female individuals. Individual data points and trend lines fitted using Locally Estimated Scatterplot Smoothing (LOESS) and 95% confidence intervals are plotted. Pearson, Spearman, and Kendall correlations between aforementioned physical parameters and age, for every sex, were calculated. Wilcoxon rank-sum test was used to compare the age distribution of individuals that made mistakes, or did not, during a narrow walk exercise. Data are plotted using ‘geom_boxplot’, with a line at the median, box height representing interquartile range from quartile 3 to 1, upper line extending up to the 75^th^ percentile, and bottom to the 25^th^ percentile. Points above the 75^th^ or below the 25^th^ percentile indicate outliers. Data and extended statistical analyses of these data are available in supplementary tables.

### Data accessibility

Raw and processed sequencing files for nRNA-seq of astrocytes in the cerebellum of aged mice fed control or PLX3397-formulated diets are available under GSE309433. Table containing gene normalized counts, fold changes and differential expression analysis is included in Supplementary tables. Raw and processed sequencing files for single-nucleus RNA-seq of astrocytes in cortex and cerebellum of adult and aged mice are available under GSE309431. Raw and processed sequencing files for ATAC-seq of astrocytes in cortex and cerebellum of adult and aged mice are available under GSE309431. All source data required for interpretation of this manuscript are available in Supplementary tables. SLEAP models, behavioral videos, and MATLAB code used in beam kinematic analysis are available in <u>10.5281/zenodo.17477698.</u>

### Statistical analysis

Statistical analysis of omics datasets is explained in the respective method details section. GraphPad prism (v 10.5.0) was used for statistical analysis and graphics of non-omics data, and R software v4.4.2 and Rstudio v2023.6.2.561 for human behavioral data. A significance threshold of alpha = 0.05 was used. For comparisons between two groups, both a Welch’s *t*-test and a Mann-Whitney test were conducted, and any observed differences were reported if identified by either test. For comparisons among more than two groups 2-way ANOVA or repeated measures (RM) 2-way ANOVA followed by Sidak’s multiple comparisons test were used. Statistical testing methodology is specified in each figure legend.

